# Amino acid Mutations at the Interface of Sudan Virus VP40 alter transport and budding

**DOI:** 10.64898/2026.05.29.728849

**Authors:** Anke-Dorothee Werner, Wieland Steinchen, Christopher Veeck, Martin Schauflinger, Laura Werel, Gert Bange, Lars-Oliver Essen, Stephan Becker

## Abstract

The matrix protein VP40 of orthoebolaviruses coordinates virion release and downregulates viral RNA synthesis through distinct oligomeric states, including dimers, octamers, and filamentous assemblies. To dissect the contributions of two oligomeric interface residues, L117 and W95, in the Sudan virus (SUDV) VP40 (sVP40), we created variants carrying alanine substitutions and assessed their structural and functional properties. sVP40 L117A failed to form dimers and was predominantly monomeric showing increased structural flexibility, reduced thermal stability together with loss of plasma membrane transport, budding activity, and the ability to regulate viral RNA synthesis. VP40 W95A preserved dimerization but also exhibited increased structural flexibility and reduced thermal stability. Functionally, sVP40 W95A more strongly inhibited viral RNA synthesis and markedly enhanced budding. However, in a transcription- and replication-competent virus-like particle (trVLP) assay, trVLPs produced with sVP40 W95A induced substantially reduced reporter activity in target cells, indicating impaired particle infectivity or functionality and suggesting possible defects in minigenome packaging, entry, or early post-entry steps. These results demonstrate that mutations at key oligomerization interfaces exert distinct structural and functional effects and highlight the requirement for precise oligomerization in coordinating sVP40’s dual roles in genome regulation and virion release. By defining the contributions of L117 and W95, this study advances mechanistic understanding of sVP40 function and identifies processes that may serve as targets for antiviral intervention.

**Importance:** Sudan virus (SUDV) causes regular outbreaks in Sub-Sahara Africa with unusually high lethality rates. However, in contrast to the more often occurring Zaire ebolavirus (EBOV), no monoclonal antibodies or vaccines are available and SUDV is generally understudied. The matrix protein VP40 is responsible for the downregulation of viral genome replication and transcription as well as budding. Here, we present structural and functional characterization of the SUDV VP40 interface residues L117 and W95 and show that while both amino acids are crucial for VP40’s structural integrity, their functional effects are dramatically different ranging from complete abolishment to improving regulatory and budding activities.

## 1 Introduction

The genus *Orthoebolavirus* belongs to the family of *Filoviridae* (order *Mononegavirales)* and comprises six species: *Orthoebolavirus zairense* (Ebola virus, EBOV), *Orthoebolavirus sudanense* (Sudan virus, SUDV), *Orthoebolavirus restonense* (Reston virus, RESTV), *Orthoebolavirus taiense* (Taï Forest virus, TAFV), *Orthoebolavirus bundibugyoense* (Bundibugyo virus, BDBV) and *Orthoebolavirus bombaliense* (Bombali virus, BOMV) (*1*). EBOV and SUDV were responsible for the majority of filovirus outbreaks in the past (*2*).

THE ROD-SHAPED MORPHOLOGY OF ORTHOEBOLAVIRUSES IS CAUSED BY THE MATRIX PROTEIN VP40, WHICH ADOPTS SEVERAL OLIGOMERIC FORMS, ALL OF WHICH EXERTING DISPARATE FUNCTIONS (*3*). THE BUILDING BLOCK FOR HIGHER-ORDER OLIGOMERS OF EBOV VP40 (EVP40) IS A DIMER WHOSE MONOMERS ARE CONNECTED THROUGH AMINO ACID RESIDUES 52-65 AND 108-117 OF THE N-TERMINAL DOMAINS (NTD;

A to C). Dimerization of eVP40 is essential for its transport towards the plasma membrane, which includes the COPII vesicular transport system (*4*). At the plasma membrane, eVP40 dimers interact with phosphatidylserine which triggers the formation of filamentous eVP40 composed of chains of eVP40 dimers (*5*) connected via their C-terminal domains (CTD). Filamentous eVP40 is stabilized by residues L203, I237, M241, M305 and l307, which form a smooth and hydrophobic interface (Figure 1 A, D and E) (*6*). Budding activity of eVP40 is dependent on late domain motifs in the N-terminus of the protein. The motifs 7-PTAP-10 and 10-PPxY-13 recruit proteins of the ESCRT complex which eventually enables budding of new virions and their abscission from the plasma membrane (*7*).

**Figure 1:**
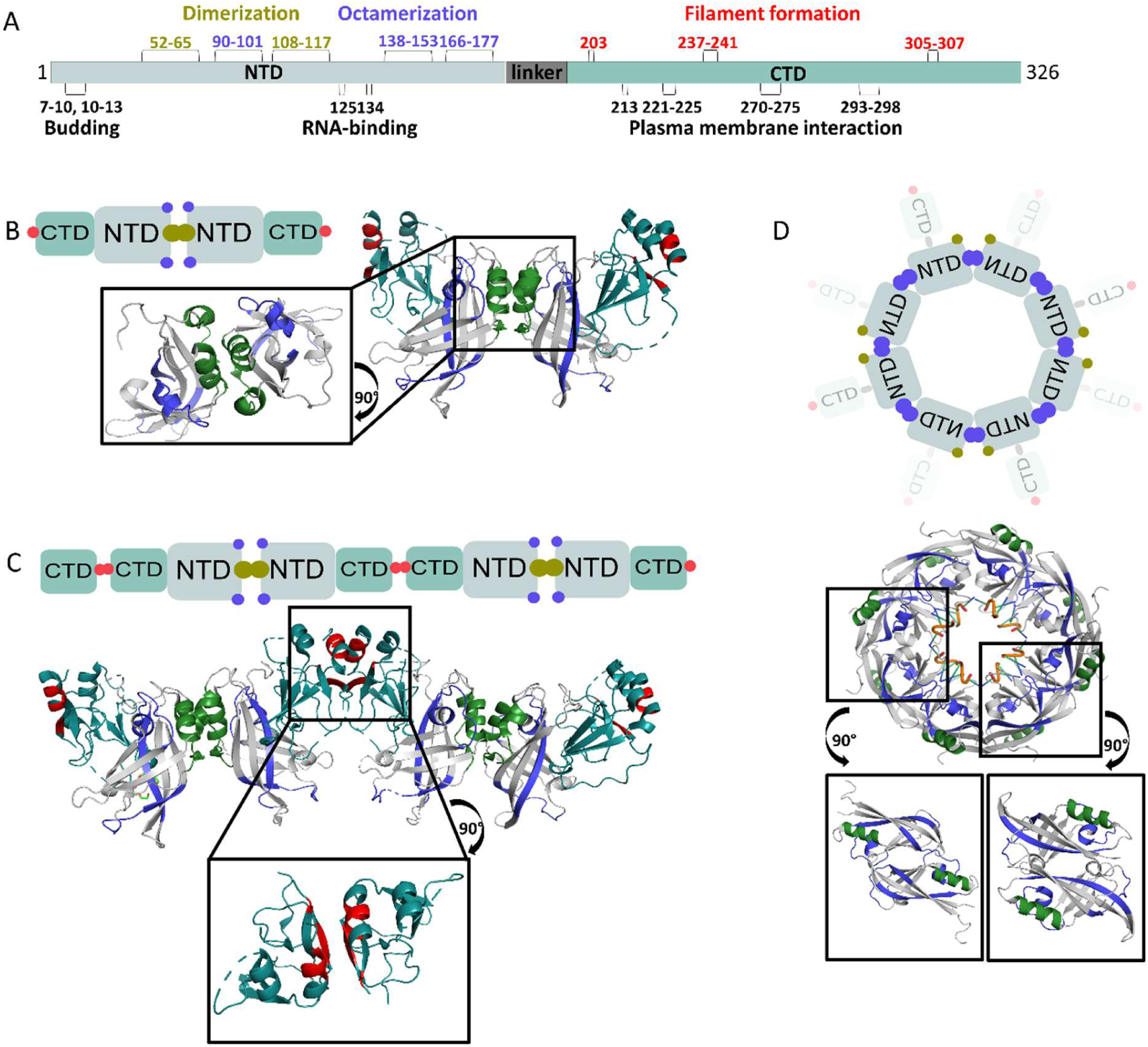
Schematic representation of sVP40 oligomer structures. A) Residues involved in homo-oligomerization interfaces are indicated above the sequence graph: dimeric interface (green), octameric interface (blue), and filamentous interface (red). Additional residues of functional interest are shown below the sequence in black. B) VP40 dimers (PDB: 4LDD) connected via the dimeric interface (green). C) VP40 filaments (PDB: 7JZJ) connected via the dimeric (green) and filamentous (red) interfaces. D) VP40 octamer (PDB: 1H2C) connected via the octameric interface (purple). For each oligomer, a close-up view of the respective interface(s) is shown below the crystal structure.

Dimeric eVP40 can be triggered to form ring-shaped octamers by interaction with RNA (*8*). The CTDs of the octamer appear to be flexible and located either above or below the eight protomers of the octamer and the NTDs are connected through different interfaces with Trp95, Glu160 and others as hot spot amino acids. Crystals of eVP40 octamers showed 5’-UGA-3’ trimeric nucleotides bound in the inside of a ring-like structure, which are connected to the residues R134 and F125 of each monomer (Figure 1 A, F and G) (*9*). In the presence of octameric eVP40 replication and transcription of the viral genome is reportedly inhibited (*10*). To broaden the knowledge about VP40 proteins of orthoebolaviruses other than EBOV, we investigated the matrix protein VP40 of SUDV (sVP40). Our objective was to functionally validate two putative hotspot amino acids, tryptophan 95 and leucine 117, located at the dimer interface of sVP40, employing a combination of structural, biochemical and functional approaches.

The investigation of SUDV proteins is of particular relevance given the recent re-emergence of SUDV as a significant public health threat, with several outbreaks reported in Central Africa (*11, 12*). These outbreaks underscore the urgent need for a better molecular understanding of SUDV, especially as no approved vaccines or antiviral therapies are currently available for this orthoebolavirus species. A detailed characterization of key viral proteins such as sVP40 may thus contribute to the development of targeted countermeasures (*13*).

## 2 Results

### 2.1 Mutation of both W95 and L117 to alanine influences the functions of sVP40 in the viral replication cycle

Based on available high-resolution structural data, residues leucine at position 117 (L117) and tryptophan at position 95 (W95) were identified as critical for VP40 oligomerization, with L117 and W95 presumed essential for dimer and octamer stability, respectively. To further evaluate their functional roles, we generated alanine substitution mutants, sVP40 L117A and sVP40 W95A.

To investigate the capacity of sVP40 to inhibit viral replication similarly to eVP40, we employed a minigenome (MG) assay (*28*). Due to the absence of a SUDV-specific minigenome system, we adapted the available EBOV system by substituting eVP40 with sVP40. As shown in Figure 2A, wild-type sVP40 (sVP40 WT) inhibited reporter gene expression in a dose-dependent manner, although it was less effective than eVP40 when equivalent plasmid amounts were used. While sVP40 W95A showed an enhanced suppression of reporter gene activity compared to sVP40 WT (Figure 2B), sVP40 L117A failed to inhibit viral genome replication and transcription, demonstrated by a substantial increase in reporter gene activity relative to sVP40 WT.

**Figure 2:**
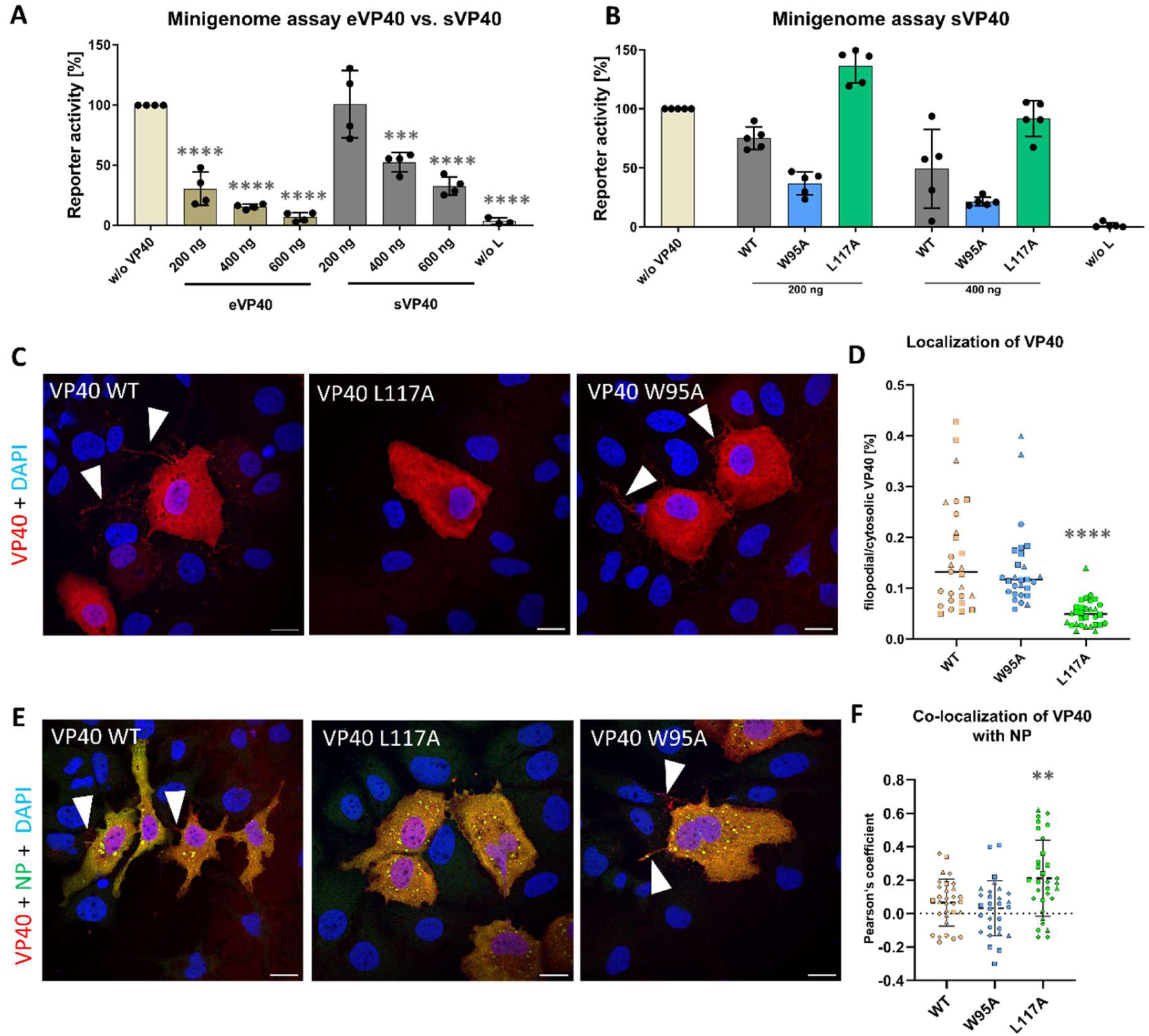
sVP40 L117A and W95A have opposing effects on VP40-regulated RNA synthesis and localization. (A–B) Effect of sVP40 WT and mutants on reporter gene activity in an EBOV-based minigenome (MG) assay. (A) Comparison of eVP40 and sVP40 WT. Asterisks indicate statistical significance (one-way ANOVA) relative to the positive control without VP40. (B) Reporter gene activity of sVP40 W95A and L117A compared with sVP40 WT. Activity was normalized to the sample without VP40 (set to 100%). (C) Representative images of immunofluorescence analyses: HuH7 cells were transfected with plasmids coding for sVP40 WT/L117A/W95A in the absence (C and D) or presence (E and F) of NP-coding plasmids. 24 h p.t., cells were fixed, and stained with protein-specific antibodies (rabbit α-sVP40 and chicken α-NP as primary antibodies and goat α-rabbit A594 and goat α-chicken A488 as secondary antibodies). Nuclei were visualized with DAPI. Confocal images were acquired using a Leica TCS SP5 microscope. sVP40-positive filaments are indicated by white arrowheads. Scale bars 20 µM. Quantification of intracellular and extracellular/filopodia-associated VP40 as well as the co-localization with NP were quantified (D and F, respectively). Asterisks indicate statistical significance (one-way ANOVA) compared with sVP40 WT. Bars represent mean ± SD of at least three independent experiments. Statistical significance: *P < 0.05, **P < 0.005, ***P < 0.0005, ****P < 0.0001.

Intracellular distribution of sVP40 WT and the interface mutants, sVP40 W95A and sVP40 L117A, was detected by immunofluorescence analysis (Figure 2C–F). As expected, sVP40 WT displayed a broadly diffuse cytoplasmic distribution with pronounced VP40-specific signal at the cell periphery, particularly in filopodia-like protrusions (white arrowheads, Figure 2C), consistent with previous observations (*14*). A similar pattern could be observed for sVP40 W95A. In contrast, the sVP40 L117A mutant showed a marked signal reduction at the plasma membrane and in filopodia-like structures (Figure 2C and D). Quantification confirmed that peripheral VP40 signal was strongly diminished, indicating impaired plasma membrane association (Figure 2D). This result suggested that the observed phenotype reflects defective trafficking and/or membrane interaction of VP40 L117A.

Upon co-expression with NP, both sVP40 WT and W95A were partially recruited to NP-induced inclusion bodies, showing moderate co-localization in the perinuclear region while still forming VP40-positive filamentous structures at the cell periphery (Figure 2E and F). In contrast, the L117A mutant exhibited a pronounced increase in co-localization with NP, accompanied by a loss of peripheral VP40 structures.

Quantitative analysis using Pearson’s (correlation) coefficient (PCC) further supports these observations. For sVP40 WT and W95A, PCC values were close to 0 (approximately 0–0.05), indicating largely random overlap between VP40 and NP signals, which is expected given the predominantly diffuse cytoplasmic distribution of VP40. In contrast, the L117A mutant showed a markedly increased PCC (approximately ∼0.3), reflecting a significant shift toward co-localization with NP-positive inclusion bodies. While a PCC of ∼0.3 may appear moderate, it represents a significant increase over random distribution in this context, particularly considering the large cytoplasmic pool of VP40. Thus, the increase in PCC for L117A suggests a biologically meaningful redistribution of VP40 rather than a minor statistical effect.

Taken together, these data suggest that mutation of L117 alters the intracellular equilibrium of VP40 localization, impairing membrane association while promoting accumulation in NP-containing inclusion bodies, thereby indicating a functional link between VP40 dimer interface integrity, intracellular trafficking and interaction with viral inclusion bodies.

Upon ectopic expression, VP40 is capable of mediating its own release into the culture supernatant in the form of virus-like particles (VLPs) (*15*). This budding activity is enhanced by co-expression of VP40 with the viral glycoprotein GP (*15, 16*). To assess whether mutations at residues L117 and W95 affect the budding capacity of sVP40, wild-type or interface mutants of sVP40 were co-expressed with GP, and both cell lysates and supernatants were analyzed for the presence of VP40 (Figure S1B and D) (*16*). Western blot analysis of the supernatants revealed a markedly increased signal for sVP40 W95A-induced VLPs in comparison to the wild-type, while only trace amounts of sVP40 L117A were detectable (Figure 3A), indicating that the L117A mutation may impair intracellular trafficking or budding efficiency, as already suggested by the immunofluorescence analysis (Figure 2C-F).

**Figure 3:**
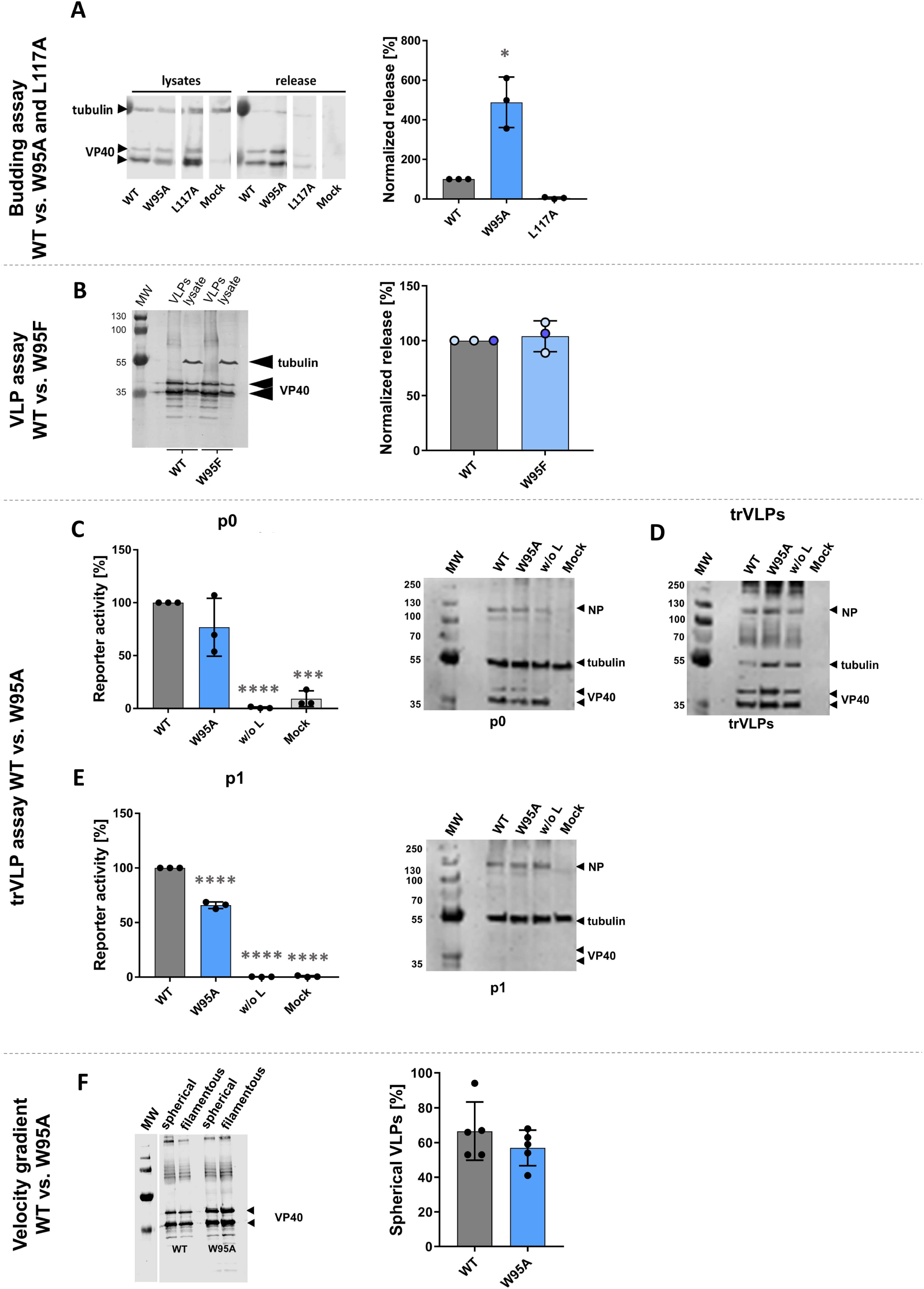
Mutation of sVP40 W95 to alanine but not phenylalanine increases budding but decreases formation of trVLPs. A) Effect of W95A and L117A mutations on VP40 release. Supernatants and cell lysates were collected 72 h post transfection (p.t.). Western blot (WB) analyses of cellular and released VP40 were performed and released VP40 was normalized to the amount of VP40 in lysates, with sVP40 WT set to 100%. All VP40-specific bands were used for quantification. B) sVP40 W95F exhibits budding activity comparable to WT levels in a budding assay (as described for A). C–E) sVP40 W95A forms less infectious transcription- and replication competent (tr)VLPs. P0 producer cells (HEK293) were transfected with pCAGGS NP, VP30, L, VP35, pANDY EBOV minigenome (3E5E), pCAGGS T7, pGL4 firefly luciferase, and either pCAGGS sVP40 WT or W95A. P0 cells were lysed 72 h p.t., and supernatants cleared of cellular debris. trVLPs were collected by ultracentrifugation (2 h at 40,000 rpm) and resuspended in 100 µL PBS_def_ 30 µL of trVLPs were analyzed by WB; 70 µL were used to infect pre-transfected P1 cells (HuH7). At 72 h post infection (p.i.), p1 cells were lysed and reporter gene activity measured. WB analyses of p0 and p1 lysates were performed using chicken α-NP, rabbit α-VP40, and mouse α-tubulin as primary antibodies, with donkey α-mouse and goat α-rabbit IRDye® 680 as secondary antibodies. F) sVP40 W95A forms more filamentous than spherical VLPs. trVLPs harvested 72 h p.t. were separated into spherical and filamentous fractions by nycodenz gradient centrifugation (15 min at 16,000 rpm). Fractions were concentrated by ultracentrifugation, resuspended, and analyzed by WB. Asterisks indicate statistical significance (one-way ANOVA) compared with sVP40 WT. Bars represent mean ± SD of at least three independent experiments. Statistical significance: *P < 0.05, **P < 0.005, ***P < 0.0005, ****P < 0.0001. MW, molecular weight ladder, given in kilo Dalton.

To evaluate the structural integrity of VLPs produced by sVP40 WT and sVP40 W95A, we performed negative staining electron microscopy (EM) (Figure S2). The morphology of the VLPs appeared highly similar between the two variants. Consistent with the Western blot results, a substantially higher number of VLPs was observed for sVP40 W95A (Figure S2). As the mutation from tryptophan to alanine represents a substantial change in side-chain size and chemical properties, we asked whether less dramatic substitutions would partially rescue the effect in functional assays. To this end, we mutated W95 to phenylalanine (W95F) which mimics the hydrophobic microenvironment more closely than W95A and performed a budding assay. Because sVP40 W95F showed a phenotype that was identical to sVP40 WT (Figure 3B), we conclude that this highly hydrophobic environment determined by a large hydrophobic sidechain is needed for protein stability and function, which is maintained upon W-to-F conversion but not W-to-A. As the phenotype of sVP40 W95F was indistinguishable from sVP40 WT, we did not pursue this mutant further.

We next employed a transcription- and replication-competent (tr)VLP assay to determine whether sVP40 W95A-induced VLPs were capable of entering target cells and initiating reporter gene expression (Figure S1) (*18*). Reporter gene analysis of the transfected producer cells (p0; Figure 3C) indicated that sVP40 W95A exerted a stronger inhibitory effect on minigenome transcription and replication compared to sVP40 WT, although the difference was not statistically significant. The expression of sVP40 WT and sVP40 W95A in both producer cells and trVLPs was confirmed by Western blot (Figure 3C and D). In cells infected with trVLPs generated from sVP40 W95A-expressing producer cells, renilla luciferase activity indicating successful trVLP infection and replication was significantly reduced compared to cells infected with trVLPs from sVP40 WT expressing cells (p1; Figure 3E). A trVLP assay in HuH7 cells yielded similar results compared to HEK293 cells (Figure S3).

Since released trVLPs can adopt both spherical and filamentous morphologies – with filamentous forms more closely resembling authentic SUDV particles and exhibiting higher infectivity in the trVLP assay (*18*) – we investigated whether the W95 mutation influenced particle morphology. To this end, velocity gradient-based fractionation of cell culture supernatants was performed to separate spherical from filamentous VLPs, as previously described (*18, 19*) (Figure S1B and F). Western blot analysis of the gradient fractions revealed that, compared with sVP40 WT, the relative abundance of sVP40 W95A was reduced in fractions enriched for spherical VLPs (Figure 3F). This shift in particle distribution indicates that the reduced functionality of sVP40 W95A-derived trVLPs cannot be explained by a loss of filamentous particle formation. Rather, the data suggest that W95A promotes efficient release of VP40-containing particles but may alter other parameters that determine trVLP functionality, such as particle composition, genome incorporation or entry competence. Thus, increased budding efficiency and increased functional infectivity of released particles are not necessarily coupled in the case of sVP40 W95A.

### 2.2 L117A, but not W95A, disrupts homo-oligomerization of sVP40

To determine whether the L117A or W95A mutation affects homo-oligomerization of sVP40, size-exclusion chromatography (SEC) was performed with purified, *E. coli*-expressed sVP40_Δ43_ proteins. In all constructs (wild-type sVP40_Δ43_, sVP40_Δ43_ W95A, and sVP40_Δ43_ L117A) the N-terminal 43 amino acids (aa) of sVP40 were deleted to improve expression and down-stream crystallization.

Wild-type sVP40_Δ43_ predominantly formed dimers, with only minor amounts of octameric species detected (Figure 4A). Similarly, the sVP40_Δ43_ W95A mutant primarily eluted as dimer (Figure 4B). In contrast, the sVP40_Δ43_ L117A mutant was found mainly as a monomer, with only trace amounts of higher-order oligomers (Figure 4C and Figure S4). These results suggest that mutation of L117, but not W95, disrupts the ability of sVP40 to form dimers and higher-order oligomers.

**Figure 4:**
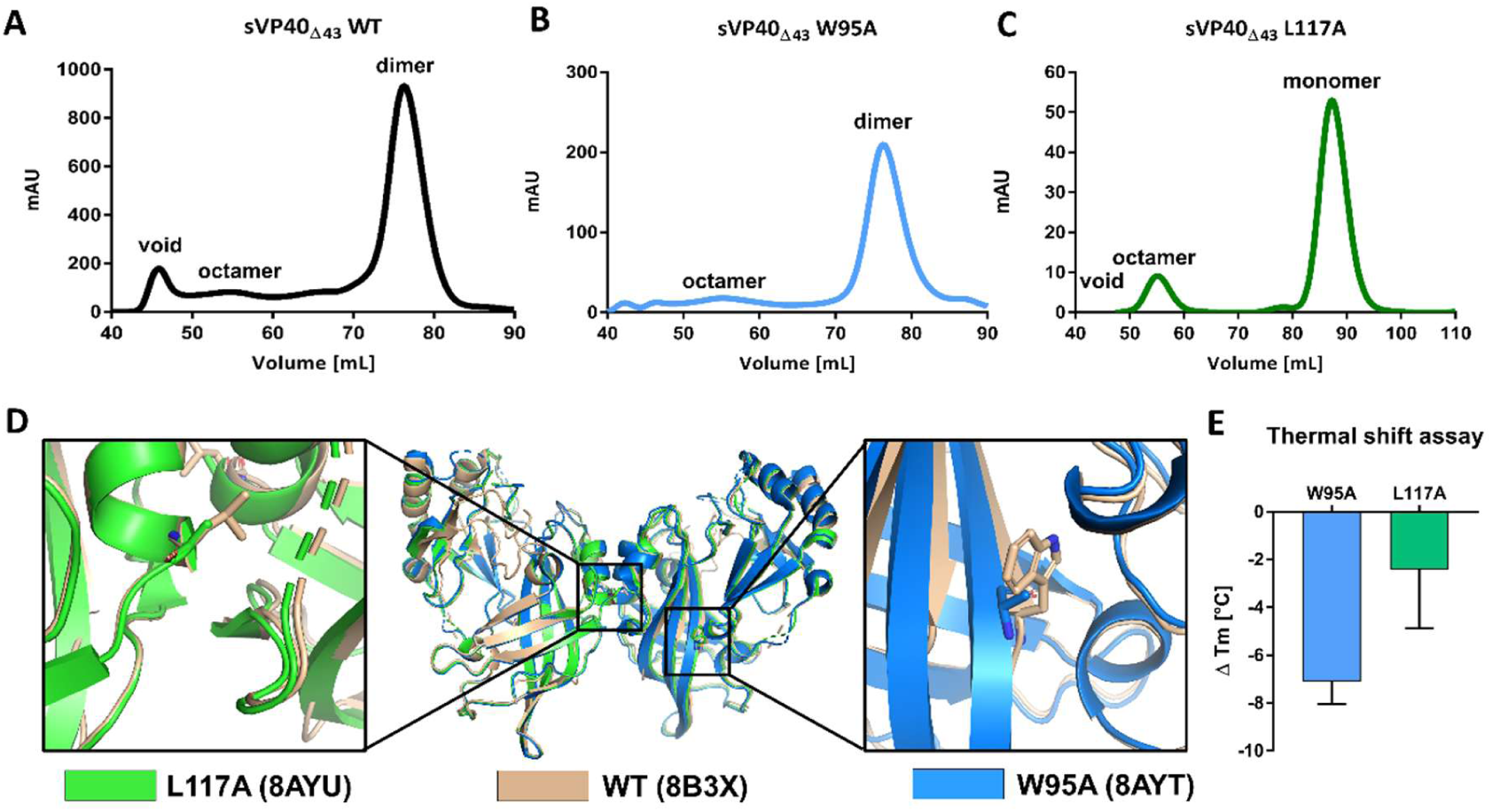
Only sVP40 L117A shows impaired oligomerization, whereas both mutants display increased flexibility and reduced thermal stability. (A–C) Size-exclusion chromatography elution profiles of (A) sVP40_Δ43_ WT (black), (B) sVP40_Δ43_ W95A (blue), and (C) sVP40_Δ43_ L117A (green), obtained using a HiLoad 16/60 column (Cytiva). (D) Structural alignment of sVP40_Δ43_ W95A and L117A crystal structures with sVP40_Δ43_ WT. Superposition of sVP40_Δ43_ WT (wheat) with W95A (PDB: 8AYT, blue) and L117A (PDB: 8AYU, green), with close-up views of the mutated residues: L117 (left panel) and W95 (right panel). (E) Thermal stability of sVP40_Δ43_ mutants. Denaturation temperatures (Tm) of sVP40_Δ43_ W95A (blue) and L117A (green) were compared with sVP40_Δ43_ WT (Tm = 49.3 ± 2.1 °C). Bars represent mean ± SD of three independent experiments.

### 2.3 sVP40Δ43 W95A and L117A are less heat-stable than sVP40Δ43 WT

To further assess the impact of the W95A and L117A mutations on the stability of sVP40, thermal shift assays (TSA) were performed. As shown in Figure 4E, the melting temperature (Tm) of wild-type sVP40_Δ43_ was 49.3 ± 2.1 °C, while the Tm values for the W95A and L117A mutants were significantly reduced to 42.2 ± 1.2 °C and 46.9 ± 0.4 °C, respectively. These results indicated that both point mutations compromise protein stability, with the W95A mutation having a more pronounced effect.

### 2.4 Hot spot mutants exhibit the same crystal packing as sVP40Δ43 WT

SEC fractions corresponding to the dimeric form of sVP40_Δ43_ and sVP40_Δ43_ W95A as well as the monomeric form of sVP40_Δ43_ L117A were collected and used to set up crystallization screens in order to investigate potential structural changes introduced by the W95A and L117A mutations. Multiple datasets were obtained from crystals grown under varying conditions, and the crystals exhibiting the highest diffraction quality were selected for further analysis. The published structure of dimeric sVP40_Δ43_ WT (PDB ID: 4LD8) (*3*) served as a template for molecular replacement. The 4LD8 model was confidently placed into the X-ray-derived electron density maps and refined using the PHENIX refinement suite (*20*) and COOT (*21*). Crystal structures of sVP40_Δ43_ W95A and sVP40_Δ43_ L117A both belonged to space group C121 and contained a single protomer in the asymmetric unit, with comparable unit cell dimensions. Experimental details for the selected crystals are summarized in Table 1.

**Table 1:**
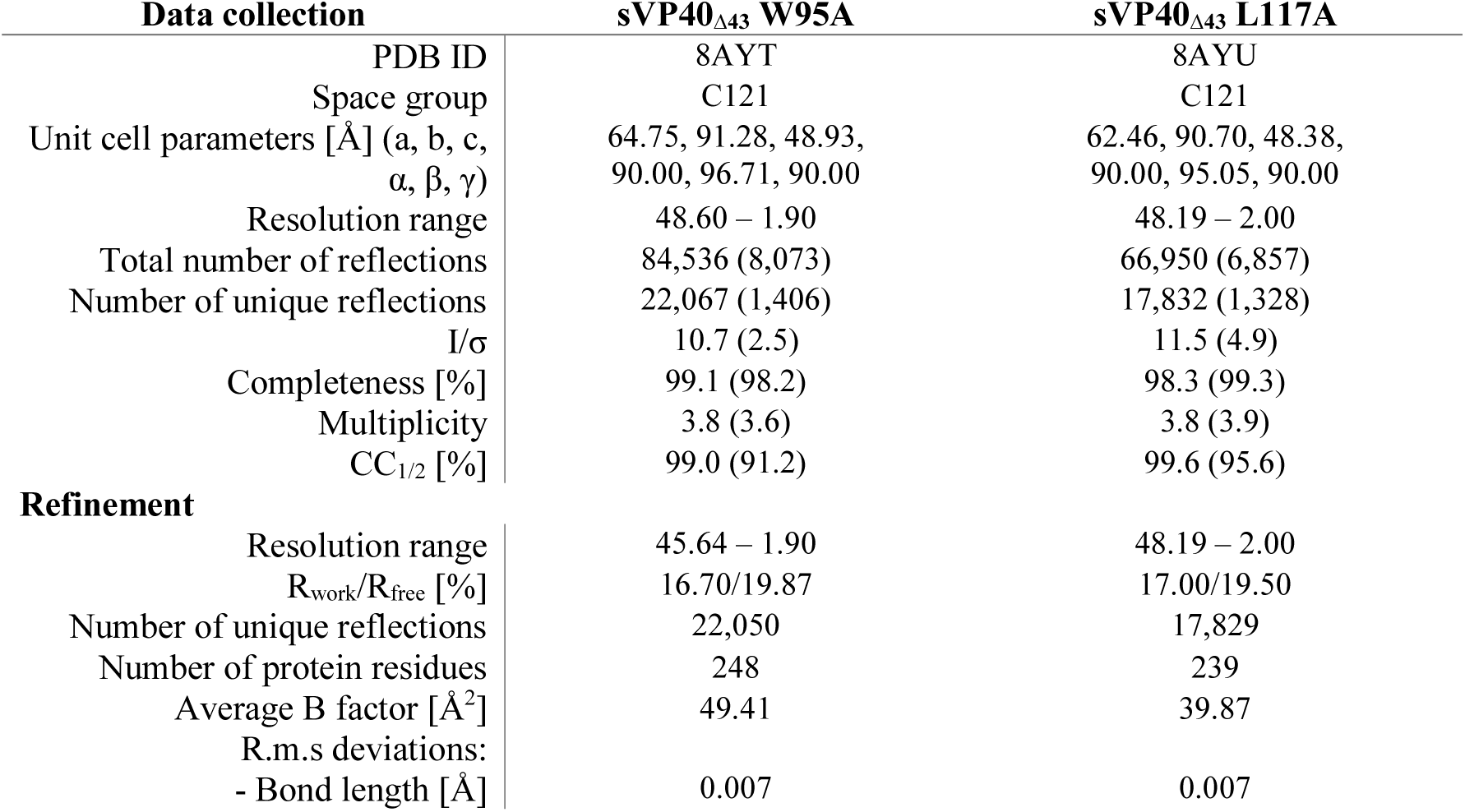
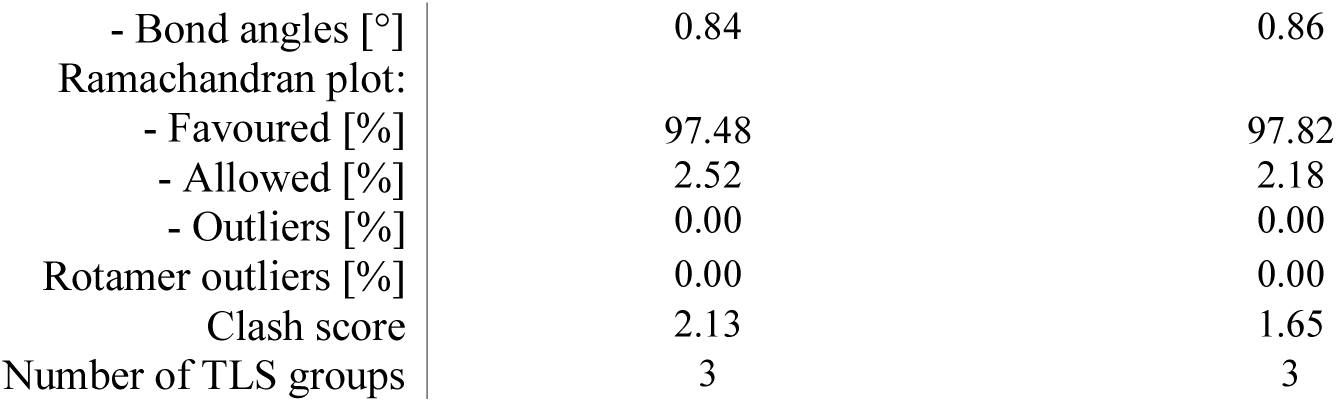
Data collection and refinement statistics of sVP40_Δ43_ W95A and L117A. Values for the outer shell are given in parentheses.

| **Data collection** |  | **sVP40<sub>Δ43</sub> W95A** | **sVP40<sub>Δ43</sub> L117A** |
| --- | --- | --- | --- |
| PDB ID |  | 8AYT | 8AYU |
| Space group |  | C121 | C121 |
| Unit cell parameters [Å] (a, b, c, α, β, γ) |  | 64.75, 91.28, 48.93, 90.00, 96.71, 90.00 | 62.46, 90.70, 48.38, 90.00, 95.05, 90.00 |
| Resolution range |  | 48.60 – 1.90 | 48.19 – 2.00 |
| Total number of reflections |  | 84,536 (8,073) | 66,950 (6,857) |
| Number of unique reflections |  | 22,067 (1,406) | 17,832 (1,328) |
| I/σ |  | 10.7 (2.5) | 11.5 (4.9) |
| Completeness [%] |  | 99.1 (98.2) | 98.3 (99.3) |
| Multiplicity |  | 3.8 (3.6) | 3.8 (3.9) |
| CC<sub>1/2</sub> [%] |  | 99.0 (91.2) | 99.6 (95.6) |
| **Refinement** |  |  |  |
| Resolution range |  | 45.64 – 1.90 | 48.19 – 2.00 |
| R<sub>work</sub>/R<sub>free</sub> [%] |  | 16.70/19.87 | 17.00/19.50 |
| Number of unique reflections |  | 22,050 | 17,829 |
| Number of protein residues |  | 248 | 239 |
| Average B factor [Å<sup>2</sup>] |  | 49.41 | 39.87 |
| R.m.s deviations: |  |  |  |
| - Bond length [Å] |  | 0.007 | 0.007 |

|  |  |  |
| --- | --- | --- |
| - Bond angles [°] | 0.84 | 0.86 |
| Ramachandran plot: |  |  |
| - Favoured [%] | 97.48 | 97.82 |
| - Allowed [%] | 2.52 | 2.18 |
| - Outliers [%] | 0.00 | 0.00 |
| Rotamer outliers [%] | 0.00 | 0.00 |
| Clash score | 2.13 | 1.65 |
| Number of TLS groups | 3 | 3 |

Both mutants crystallized in a dimeric arrangement, with packing closely resembling that of the wild-type sVP40, as reflected by the low root-mean-square deviation (RMSD) values: 0.417 Å for sVP40_Δ43_ WT vs. W95A and 0.287 Å for sVP40_Δ43_ WT vs. L117A (Figure 4D). While dimer formation was expected for sVP40_Δ43_ W95A, it was notable that the sVP40_Δ43_ L117A mutant, which was predominantly monomeric in solution, also adopted a dimeric arrangement in the crystalline state. This suggests that dimeric packing represents a particularly stable configuration under crystallization conditions, even for mutants with impaired oligomerization in solution. The crystallographic B-factors serve as indicators of atomic mobility and structural flexibility. Both mutant structures exhibited elevated B-factors compared to the wild-type, with particularly high values observed for the sVP40_Δ43_ W95A mutant (Table 1, Figure S5), supporting the conclusion that these mutations increase the dynamic disorder of the protein. This result is also reflected in the reduced thermal stability of both mutants (Figure 4E).

### 2.5 HDX-MS reveals major structural implications in solution for hot spot mutants

The previous results suggested that conformational changes especially in sVP40_Δ43_ L117A may not be fully represented in the available crystal structures. To further assess potential conformational rearrangements in solution, we performed hydrogen–deuterium exchange mass spectrometry (HDX-MS). Initial analyses were carried out using sVP40_Δ43_ and exchange rates were plotted onto the protein structure (Figure 5). This experiment revealed that residues within the dimer interface (spanning aa 109–117) exhibited very low deuterium uptake with slow exchange kinetics, consistent with limited solvent accessibility (Figure 5, black box). Likewise, residues 73–81 and 90–99, including W95, exhibited very slow HDX due to their embedment in a ß-sheet (Figure 5, black dotted box). Overall, the HDX profile of sVP40_Δ43_ WT closely resembled that of Marburg virus VP40 (*22*) and corroborated the crystallographic data for most structural regions. The reduced deuterium uptake observed around L117 and W95 indicates that these residues are substantially buried within the protein core in solution, in contrast to what would be expected for a VP40 monomer, thereby underscoring their critical contribution to stabilizing the native conformation of VP40 in solution.

**Figure 5:**
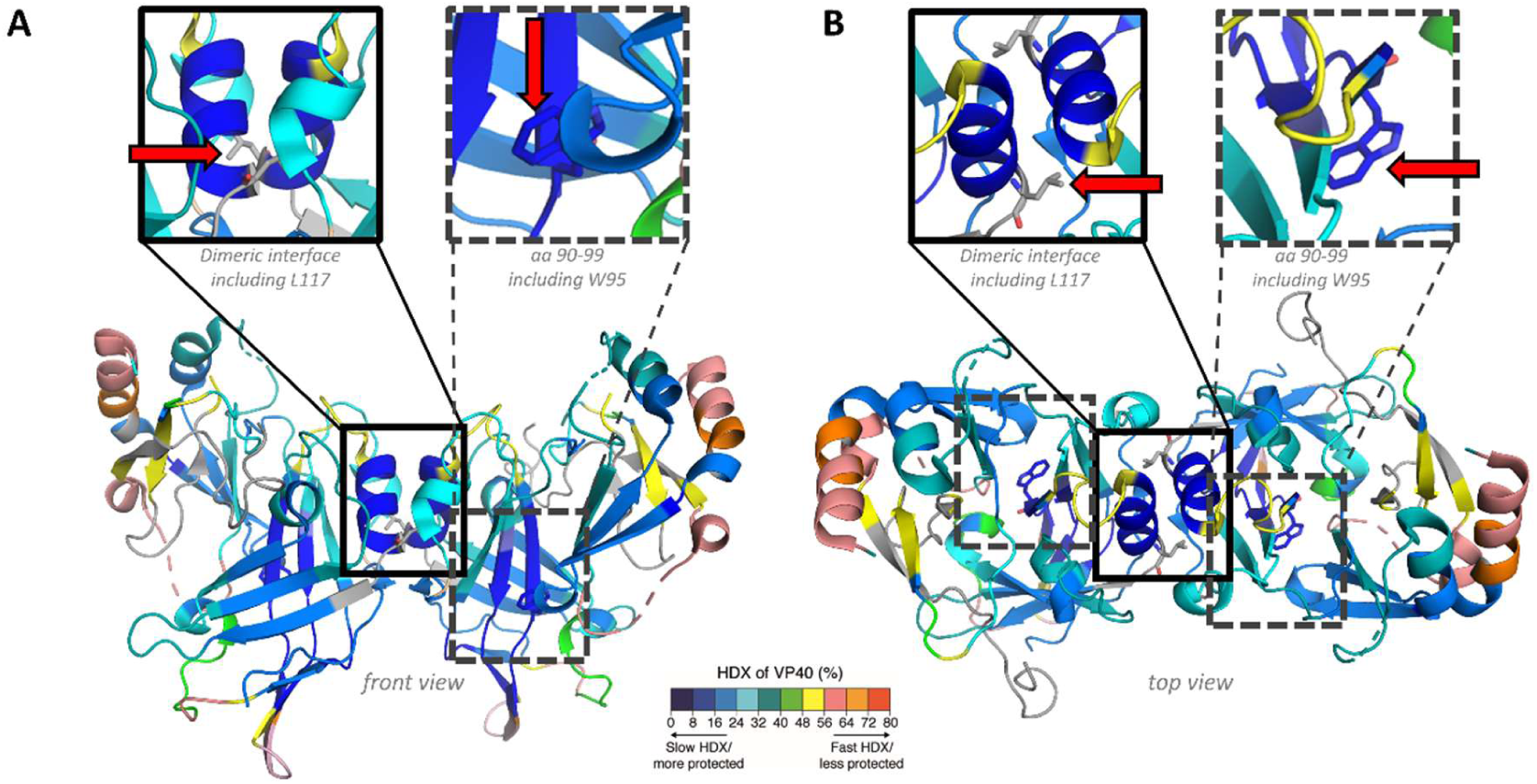
Hydrogen–deuterium exchange rates of sVP40_Δ43_ WT confirm that L117 and W95 are embedded in the protein core. sVP40_Δ43_ WT (25 µM) or mutant proteins were incubated in D₂O-containing buffer. Reactions were quenched after 10, 30, 100, 1,000, or 10,000 s, followed by pepsin digestion. The resulting peptides were ionized by ESI and analyzed by mass spectrometry. D₂O uptake of sVP40_Δ43_ WT mapped onto the dimeric crystal structure (PDB: 8B3X), Front view (A) and top view of the structure (B). Structural regions of interest are highlighted: the N-terminal dimeric interface (black solid box), the CTD-to-CTD interface of filaments (grey double-framed box), and the region containing W95 (black dotted box). Amino acids with low exchange rates are shown in blue, those with high exchange rates in red. Residues not covered in the analysis are indicated in white. Red arrows are used to indicate the position of L117 and W95.

Given the unexpected observation that sVP40_Δ43_ L117A crystallized in a dimeric conformation indistinguishable from wild-type sVP40_Δ43_, we next assessed its conformation in solution by HDX-MS. The data revealed marked differences in deuterium uptake between sVP40_Δ43_ L117A and sVP40_Δ43_ WT (Figure 6A and B and Figure S6). In particular, peptides spanning residues 108–114, and to a lesser extent 52–65 (*3*), showed increased deuterium incorporation in the mutant. These differences indicate that the dimer interface of sVP40_Δ43_ L117A is solvent-accessible and not engaged in intermolecular contacts, consistent with a mostly monomeric state in solution. Additional regions, including residues 176–180 (NTD) and 248–260 (C-terminal β-sheets), also exhibited altered exchange kinetics, suggesting that the L117A substitution influences both local secondary structure and the overall conformational dynamics of the protein. Mapping the differentially exchanging regions onto the (WT) crystal structure (Figure 6 B) further highlighted the disruption of dimer-stabilizing interactions.

**Figure 6:**
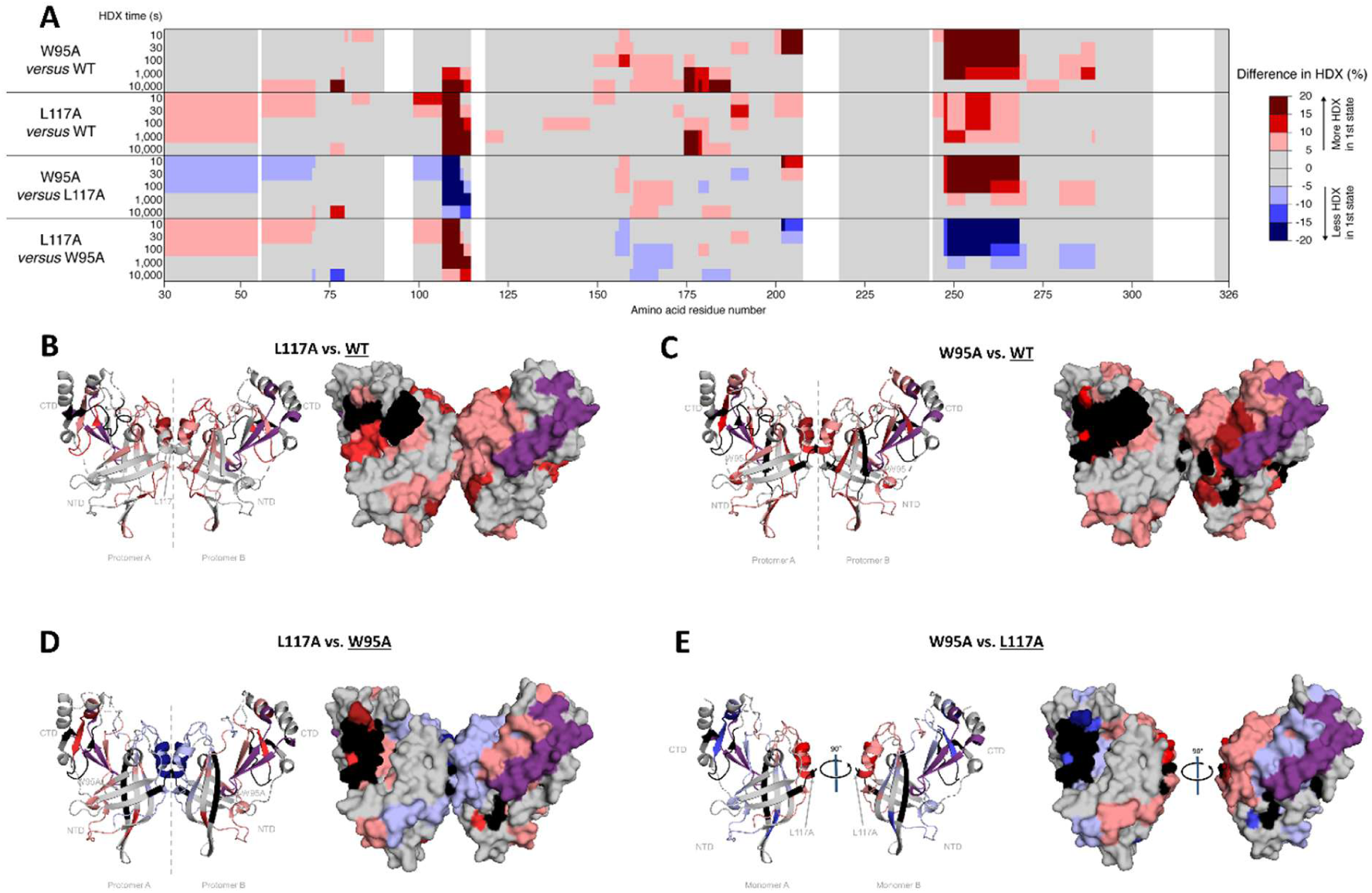
Mutation of L117 and W95 to alanine increases protein fluidity and hydrogen/deuterium (H/D) exchange. sVP40_Δ43_ wild type (25 µM) or mutant proteins were incubated in D₂O-containing buffer. Reactions were quenched after 10, 30, 100, 1,000, or 10,000 s, followed by pepsin digestion. The resulting peptides were ionized by ESI and analyzed by mass spectrometry. A) Mapping the differentially exchanging regions onto the protein sequence. B-C) Differential deuterium uptake of sVP40_Δ43_ WT vs. W95A (B) and L117A (C) mapped onto the crystal structure of WT crystal structure. D-E) Differential deuterium uptake of sVP40_Δ43_ W95A vs. L117A mapped onto the crystal structure of sVP40_Δ43_ W95A (PDB: 8AYT) (D) and vice versa (E). For clarity, only the monomeric version of L117A is depicted in panel D. Amino acids with low exchange rates are shown in blue, those with high exchange rates in red. Unchanged residues are depicted in grey, peptides exhibiting bimodality in purple, and residues not covered are indicated in black.

Taken together, the SEC data (Figure 4) and HDX-MS results (Figure 6) strongly support the conclusion that sVP40_Δ43_ L117A exists in solution either predominantly as a monomer or in rapid equilibrium between monomeric and dimeric states. Accordingly, dimer formation during intracellular expression is highly impeded and the crystal structure mostly represents a minor species available to the L117A variant in solution – presumably reflecting the most energetically favorable packing in the crystalline state.

Comparison of sVP40_Δ43_ W95A with wild-type sVP40_Δ43_ WT also revealed substantial differences in deuterium uptake (Figure 6A and C and Figure S6). Many of the same peptides affected in the L117A mutant also showed altered exchange in W95A, pointing to a destabilization, but not complete disruption, of the dimer interface. The strongest increase in deuterium uptake was observed for residues 248–260, with additional regions of elevated exchange comprising peptides 203–208, 176–188, and segments of the dimer interface.

Although the WT vs. L117A and WT vs. W95A exchange patterns appeared similar at first glance, direct comparison of the two mutants underscored their differences, e.g., the dimer interface of sVP40_Δ43_ W95A showed reduced deuterium uptake relative to sVP40_Δ43_ L117A (Figure 6D and E).

Strikingly, both mutants displayed bimodal behaviour in the same peptide, characterized by the coexistence of two distinct isotopic envelopes in the mass spectra (Figure 6, highlighted in purple; Figure S7). Such bimodality reflects conformational heterogeneity, indicating multiple structural states differing in solvent accessibility and stability. This pattern is consistent with the presence of more than one oligomeric species, e.g. due to rapid monomer-dimer equilibrium or partial unfolding under native conditions.

A comprehensive overview of all HDX-MS results is provided in Table S1.

Together, these findings reveal that the two interface residues analyzed here affect sVP40 function through distinct mechanisms. L117 is essential for maintaining the dimeric state of sVP40, and its mutation to alanine results in a broadly loss-of-function phenotype affecting intracellular distribution, budding, and regulation of viral RNA synthesis. In contrast, W95 is not required for dimer formation per se, but appears to contribute to the conformational stability and dynamic organization of the dimer. Consequently, the W95A mutation does not abolish VP40 activity but rather shifts its functional output, enhancing budding and inhibition of viral RNA synthesis while reducing the functional activity of released trVLPs.

## 3 Discussion

This study identified two residues, L117 and W95, at the oligomerization interface of Sudan virus VP40 as critical modulators of structural integrity and function. Substitution of L117 with alanine disrupted dimerization, rendering VP40 monomeric in solution and abolishing its ability to regulate RNA synthesis or drive budding. These results are in line with the impact of the homologous amino acid in eVP40 (*3*). Interestingly, in the crystal packing, sVP40_Δ43_ L117A was present as dimer and almost indistinguishable from the 3D structure of sVP40_Δ43_ (Figure 4D). Similar discrepancies between crystallographic and solution states have been reported previously, e.g., for the RdRp domain of nsp4 from Sindbis virus, which crystallized as a dimer but was shown to be monomeric in solution (*23*), for the bacterial RNA-modifying enzyme Tgt (*24*), and for calmodulin, which displayed highly rigid regions in the crystal structure that were revealed to be flexible by HDX-MS (*25*).

In contrast, substitution of W95 with alanine destabilized the protein while maintaining dimerization. The sVP40 W95A variant exhibited markedly increased budding and stronger inhibition of RNA synthesis, yet trVLPs produced with this mutant induced reduced reporter activity in P1 cells. This discrepancy is noteworthy, as the enhanced budding would normally be expected to increase the functional activity of the released particles. One possible explanation is that the stronger suppression of viral RNA synthesis in producer cells limits the amount of minigenome available for incorporation into budding particles, resulting in the release of VLPs with reduced genome content and consequently lower reporter gene expression in target cells. However, the present data do not distinguish between impaired minigenome packaging, altered particle composition, reduced entry competence, or defects during early post-entry steps. Thus, the reduced reporter activity of sVP40 W95A-derived trVLPs should be interpreted as evidence for impaired particle functionality rather than as proof for a single mechanistic defect.

W95 represents a large and hydrophobic side chain which is apparently decisive for overall protein folding in solution with strong functional implications as discussed above. Interestingly, eVP40 W95A slightly reduced octamerization but its inhibitory function on viral RNA synthesis and budding activity was unchanged (*3, 9, 10*). A comparison of the published dimeric structures of eVP40 and sVP40 WT revealed that the orientation of the W95 side chain is highly conserved (Figure S8). Its aromatic nitrogen atom forms H-bonds to the backbone oxygen atom of K256, thereby stabilizing the loop between two β-sheets. Disruption of this interaction by mutating the W to A could possibly result in a destabilization of the NTD-CTD integrity, which in turn could have implications for the formation of higher-order oligomers. Given the high crystallographic similarity between the eVP40 and sVP40 structures as well as the high stability of the dimeric packing (as also observed for the here presented mutants), differences between the two proteins are therefore unlikely to manifest in the crystalline state but rather in solution. The difference between sVP40 and eVP40 is notable because the local structural environment of W95 appears highly conserved in the available crystal structures. It suggests that even highly similar VP40 architectures may differ in their conformational equilibria, dynamic stability, or functional tolerance toward mutations at corresponding interface residues. Therefore, conclusions drawn from eVP40 cannot necessarily be transferred directly to sVP40, particularly when mutations affect buried residues that contribute to long-range structural stability. This underscores the importance of analyzing SUDV VP40 directly and supports the view that species-specific differences in VP40 dynamics may influence the balance between RNA synthesis regulation, particle formation, and particle functionality.

Our studies involving sVP40 W95F suggest that not the tryptophan side chain itself but rather its contribution to the local hydrophobic environment is decisive. Removing this hydrophobic force apparently also reflects on remote structural areas and has implications for stability, conformation and function. This supports our hypothesis, that the effects observed in cell culture experiments cannot solely be attributed to impaired interaction with other viral or host proteins. Indeed, our biochemical analyses were performed in the absence of other (viral) proteins and showed that the W95A mutation intrinsically alters the biochemical properties of the protein.

In summary, our study shows that the functional output of sVP40 is not determined alone by the presence or absence of defined oligomeric states, but also by the stability and dynamics with which these states are maintained. Whereas L117 is required to stabilize the dimeric building block that enables intracellular trafficking and budding, W95 preserves dimer formation but appears to affect the conformational equilibrium of VP40, thereby uncoupling particle release from productive trVLP function. These findings identify VP40 interface stability as a critical determinant of the balance between regulation of viral RNA synthesis and virion morphogenesis, and they highlight oligomeric interfaces and their dynamic solution-state properties as attractive targets for antiviral intervention.

## 4 Methods

### 4.1 Cells and plasmids

HEK293 (human embryonic kidney) and VeroE6 (African green monkey kidney) cells were maintained in Dulbecco’s modified Eagle’s medium (DMEM, Invitrogen) supplemented with 10% fetal calf serum (FCS), 2 mM L-glutamine, 100 U/ml penicillin and 100 μg/ml streptomycin and grown at 37 °C and 5% CO_2_. All plasmids used in this study (pCAGGS NP, pCAGGS VP35, pCAGGS VP30, pCAGGS L, pCAGGS VP40 and pET46 EK/LIC sVP40_Δ43_, and pCAGGS GP; the EBOV-specific minigenome (pANDY 3E5E-luc), and pCAGGS T7 polymerase) were already described previously (*3, 14, 26, 27*). Mutagenesis of sVP40 in pCAGGS or pET46 EK/LIC was performed using the Quickchange Multi Site-directed Mutagenesis Kit (Agilent) according to manufacturer’s instructions. The primer sequences are as follows: ctcaacaacagcagcaattatggccgcatcttatacgatcacc, gtcctcatgaaacaaatccctattgcgttgccactc-ggaa and ccaaagtcctcatgaaacaaatccctattttcttgccactcggaattg for L117A, W95A and W95F, respectively.

### 4.2 Western blot analysis

Protein samples were applied on an SDS-polyacrylamide gel and transferred to nitrocellulose membranes (0.45 µM, Amersham). Protein-specific antibody staining was performed using primary antibodies diluted 1:1,000 in 1% milk and 0.1% (v/v) Tween-20 (in PBS_def_): rabbit-α-sVP40 (Sino Biological), chicken α-NP and mouse α-tubulin as a cellular control. Secondary antibodies were diluted 1:5,000 in the same buffer: goat α-mouse IRDye® 800, goat α-mouse IRDye® 680, goat α-rabbit IRDye® 680, donkey α-mouse IRDye® 680 and donkey α-chicken IRDye® 680 using the Odyssey® CLx imaging system.

### 4.3 Minigenome assay

8x10^5^ HEK293 in 6 well plates (50% confluency) were treated as previously described (*14, 28*). In short, 1000 ng pCAGGS L, 125 ng pCAGGS VP35, 100 ng pCAGGS VP30, 125 ng pCAGGS NP, 250 ng pCAGGS T7, 250 ng pANDY 3E5E and 100 ng pGL4 Firefly luciferase. For the initial comparisons between eVP40 and sVP40, 200 ng, 400 ng or 600 ng were added to the transfection mix, respectively. To compare sVP40 WT with the different mutants, 200 ng or 400 ng of sVP40-coding plasmid were added. Differences in the absolute amount of transfected plasmid DNA were compensated for by the addition of empty pCAGGS vector. 48 h p.t., cells were washed with PBS_def_, the cells resuspended in 1x Lysis Juice and frozen at -20 °C. After thawing, the cellular debris was spun down for 10 min at 4 °C and 13,200 rpm. The supernatants were then used for the measurement of reporter gene activity using the Beetle-Juice and Renilla-Juice BIG KITs (PJK) and measured with a CentroLB 960 luminometer (Berthold Technologies) as well as for Western blot analysis. Firefly luciferase signals were used to normalize for transfection efficiency.

### 4.4 Immunofluorescence analysis

#### Transfection and staining

2x10^5^ HuH7 cells were seeded onto cover glasses and transfected with 1000 ng pCAGGS sVP40 WT/L117A/W95A (and 500 ng pCAGGS NP for co-expressions) using 3 µL TransIT® (Mirus) per µg transfected DNA at 50% confluency. 24 h post transfection (p.t.), cells were fixed with 4% PFA for 30 min, treated for 5 min with 100 mM glycine in PBS_def_ and permeabilized using 0,1% Triton X-100 in PBS_def_ before incubation in blocking buffer (2% BSA, 5% (v/v) glycerol, 0.2% (v/v) Tween-20, 0.05% (w/v) Sodium azide in PBS_def_) for 10 min.

Protein-specific staining was performed with rabbit α-sVP40 (1:500, Sino Biological) and/or chicken α-NP (1:200), as well as goat anti-Rabbit A594 (1:400) and goat anti-chicken A488 (1:500) as secondary antibodies, together with DAPI staining.

#### Confocal Microscopy and Quantitative Colocalization Analysis

Cells were imaged using a Leica STELLARIS 8 confocal microscope (Leica Microsystems) equipped with a 63x oil immersion objective (NA 1.4). For quantitative colocalization analysis, multichannel images were processed using an automated macro-script in Fiji (ImageJ version 1.54p). To ensure high-fidelity quantification within specific cellular compartments, regions of interest (ROIs) were manually defined around cell borders. To minimize potential signal noise and non-specific cytoplasmic background, a rolling ball background subtraction (radius = 50 pixels) was applied locally within each defined ROI for both VP40 (channel 1) and NP (channel 2) channels prior to analysis. Colocalization was quantified using the Coloc 2 plugin. Pearson’s correlation coefficients (PCC) were calculated to assess the degree of overlap between the two signals. To account for potential background bias, the analysis focused on PCC values above the autothreshold, determined by the Costes’ regression method. To validate the robustness of the colocalization, the analysis included 10 Costes’ cost randomizations. To ensure statistical robustness, 30–40 single-cell analyses were performed over three to four independent experiments, with all resulting coefficients exported for comparative evaluation.

#### Image Processing and Signal Quantification

Cells were imaged using a Leica STELLARIS 8 confocal microscope (Leica Microsystems) equipped with a 63x oil immersion objective (NA 1.4). To ensure objective quantification of protein distribution, raw image data (LIF format) were processed using ImageJ/Fiji (ImageJ version 1.54p). Statistical noise filtering was performed using a conservative thresholding strategy: only pixels with intensities exceeding the background mean (μ_bg_) by at least two standard deviations (σ_bg_) were considered as specific signal (Limit of Quantification). The total corrected integrated density (𝑅𝑎𝑤𝐼𝑛𝑡𝐷𝑒𝑛_all_) and the intracellular integrated density (𝑅𝑎𝑤𝐼𝑛𝑡𝐷𝑒𝑛_cell_) were measured using the *limit to threshold* function. The extracellular signal fraction (𝑅𝑎𝑤𝐼𝑛𝑡𝐷𝑒𝑛_extra_) as calculated by subtracting the intracellular signal from the total image signal:

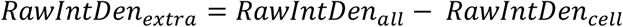

Any resulting negative values, where the signal was statistically indistinguishable from background noise, were clipped to zero to prevent mathematical artifacts. Finally, the distribution ratio of extracellular to intracellular fluorescence was calculated:

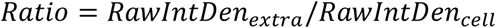

To ensure statistical robustness, 30–40 single-cell analyses were performed over three independent experiments, with all resulting factors exported for comparative evaluation.

### 4.5 Infectious and non-infectious VLP assay

For non-infectious VLP, 500 ng of sVP40 WT or L117A/W95A/W95F mutant were transfected into 8x10^5^ HEK293 cells along with 500 ng GP to increase yields (*16*). Media was changed 4 h p.t. to 3 mL DMEM containing 3% FCS. 72 h p.t. cell culture medium was collected, cell debris removed via centrifugation for 10 min at 2,500 rpm and the supernatants subjected to ultra-centrifugation for 2 h at 4 °C and 40,000 rpm using a 20% sucrose cushion and resuspended in 1x SDS sample buffer. Cells were washed once with PBS_def_, resuspended in 1x Lysis Juice (Promega) and frozen at -20 °C. After thawing, the samples were centrifuged for 10 min at 4 °C and 13,200 rpm and the supernatants were used for Western Blot analysis together with the resuspended VLPs.

Transcription and replication competent VLPs (trVLPs) were generated by transfecting 30 ng pCAGGS VP24, 250 ng pCAGGS GP, 1000 ng pCAGGS L, 125 ng pCAGGS VP35, 100 ng pCAGGS VP30, 125 ng pCAGGS NP, 250 ng pCAGGS T7, 250 ng pANDY 3E5E and 100 ng pGL4 Firefly luciferase and 500 ng pCAGGS sVP40 WT/W95A per well of HEK293 (or 2x10^5^ HuH7) cells (producer cells, p0), seeded to 50% confluency in a 6 well dish. The supernatant of three wells was pooled and pelleted by ultracentrifugation for 2 h at 40,000 rpm using a SW41 rotor (Beckman Coulter) with a 20% sucrose cushion. 100 µL PBS_def_ was used to resuspend the trVLPs, which were either used for the infection of HuH7 cells (p1, seeded to 50% confluency in a 6 well dish) or for Western blot analysis.

P1 cells (HuH7 cells, seeded to 50% confluency in a 6 well dish) were transfected with 1000 ng pCAGGS L, 125 ng pCAGGS VP35, 100 ng pCAGGS VP30 and 125 ng pCAGGS NP. 4 h p.t., the media was aspirated and replaced with 415 µL DMEM without FCS and 85 µL of the pelleted trVLP fractions. Cells were then incubated for 1 h and then 2 ml fresh DMEM containing 3% FCS were added. P1 cells were harvested 3 d p.t. as described above. Renilla signals were measured in both p0 and p1 cells (see section 4.3) and Western blot analysis was performed using samples from p0 and p1 lysates as well as the remaining 15 µL of the resuspended trVLPs.

### 4.6 Velocity gradient

The velocity gradient was performed in principle as described previously (*29*). Briefly, six wells of a 6well plate HEK293 at 50% confluency were transfected with plasmids coding for the trVLP set, as described in section 4.5. 72 h p.t., the supernatants were collected, cleared from cellular debris via low-speed centrifugation and trVLPs were collected using ultracentrifugation for 3 h at 36,000 rpm. Pellets containing the trVLPs were resuspended in 730 µL TNE buffer and applied on top of a nycodenz gradient (730 µL 30%, 490 µL 20%, 490 µL 15%, 490 µL 10%, 490 µL 7.5%, 490 µL 5%, 490 µL 2.5%, trVLP solution from bottom to top) and centrifuged for 15 min at 16.000 rpm using a SW60 rotor (Beckman Coulter). The top three fractions (fractions 1-3, 490 µL each) fractions were pooled, as were the following fractions (fractions 4-6, 490 µL each). Each pool was subjected to a third round of ultracentrifugation for 2 h at 40,000 rpm using a TLA rotor (Beckman Coulter). Pellets containing either the pooled fractions 1-3 or 4-6 were resuspended in PBS_def_ and analysed via western blot.

### 4.7 Expression, purification and crystallization of recombinant sVP40

Expression and purification were performed as described previously (*14, 30*). Briefly, the truncated *vp40* (lacking the first 43 amino acids, sVP40_Δ43_) gene was cloned into pET46 and transformed into Rosetta2 cells. The recombinant sVP40 proteins were expressed overnight at 22 °C in LB medium, induced with 0.5 mM IPTG. After expression of the proteins for 16 h at 22 °C, the bacteria were harvested and pelleted via low-speed centrifugation at 2,500 rpm. The pellets were then resuspended in 25 mM Tris, 300 mM NaCl, pH 8, 10 mM imidazole, lysed using a microfluidizer and centrifuged for 1 h at 20,000 rpm. The cleared supernatant was then incubated overnight with 1 ml Ni-NTA slurry (QIAGEN) per L expression culture. Next, the beads were washed extensively with 25 mM Tris, 300 mM NaCl, pH 8, 20 mM imidazole and, finally, the sVP40 was eluted using 25 mM Tris, 300 mM NaCl, pH 8, 250 mM imidazole. To determine the oligomeric state the recombinant protein, the solution was further purified using size-exclusion chromatography using a HiLoad 16/60 or a Superdex 200 Increase (Cytiva) column. Purified, dimeric sVP40_Δ43_ W95A was crystallized using the MBC I screen in 0.5 M potassium chloride, 50 mM MOPS, 12% PEG4000 (w/v), 20% glycerol (w/v) (well F10), with a concentration 3.6 mg protein/ml. For sVP40_Δ43_ L117A, crystals were formed in 90 mM HEPES, 6.8% ethylene glycol (v/v), 15% glycerol (v/v), 17% PEG10,000 (w/v) (Cryos screen, well H11) using 7.6 mg protein/ml. Crystallization was performed by the MarXtal facility in Marburg, Germany, using a sitting drop method. Crystals were frozen in liquid nitrogen and measured at the Swiss Light Source (SLS) of the Paul Scherrer Institute, Villigen, Switzerland, beamline PX01.

### 4.8 Data processing and refinement

Diffraction data for sVP40_Δ43_ L117A and sVP40_Δ43_ W95A were processed using XDS (*31*) and Aimless by the ccp4i suite (*32*), followed by molecular replacement using PHASER as implemented in the PHENIX suite (*20*). As a search model sVP40 dimer (PDB ID 4LD8 (*3*)) was used. Refinement for each crystal structure involved iterative cycles of model building using COOT (*21, 33, 34*) and repeated refinement cycles with PHENIX. Missing residues were modelled using COOT. Water molecules were placed manually. Side chains with missing electron density were deleted from the model. Occupancies were calculated for residues showing more than one alternative side chain orientations. The quality of the final average structure was assessed by MolProbity implemented in the PHENIX suite (*35*). Coordinates and structure factors have been deposited into the Protein Data Bank under accession codes 8AYT (sVP40_Δ43_ W95A) and 8AYU (sVP40_Δ43_ L117A).

### 4.9 HDX-MS

25 µM concentrated protein solutions of sVP40_Δ43_ WT/L117A/W95A were employed. HDX-MS experiments were conducted essentially as described previously aided by a two-arm robotic autosampler (LEAP Technologies) (*36*). In brief, 7.5 μl of sVP40_Δ43_ WT/L117A/W95A solution were mixed with 67.5 μl of D_2_O-containing buffer (25 mM Tris-Cl pH 8.0, 300 mM NaCl) to initiate the exchange reaction and incubated for 10, 30, 100, 1,000 or 10,000 s at 25 °C. Afterwards, 55 μl of sample were withdrawn from the reaction and mixed with an equal volume of quench buffer (400 mM KH_2_PO_4_/H_3_PO_4_, 2 M guanidine-HCl, pH 2.2) kept cold at 1 °C, and 95 µl of the resulting mixture was injected into an ACQUITY UPLC M-Class System with HDX Technology (Waters) (*37*). Non-deuterated samples were prepared similarly by 10-fold dilution in H_2_O-containing buffer. The proteins were digested online employing a column (2 mm x 2 cm) packed with immobilized porcine pepsin at 12 °C under a constant flow (100 μl/min) of water + 0.1% (v/v) formic acid and the resulting peptic peptides trapped on an ACQUITY UPLC BEH C18 VanGuard Pre-column, 130 Å, 1.7 µm, 2.1 mm x 5 mm (Waters) that was kept at 0.5 °C. After 3 min, the trap column was placed in line with an ACQUITY UPLC BEH C18 1.7 μm 1.0 x 100 mm column (Waters), and the peptic peptides eluted at 0.5 °C with a gradient of water + 0.1% (v/v) formic acid (A) and acetonitrile + 0.1% (v/v) formic acid (B) at 60 µl/min flow rate as follows: 0-7 min/95-65% A, 7-8 min/65-15% A, 8-10 min/15% A. The eluting peptides were ionized by electrospray ionization (capillary temperature 250 °C, spray voltage 3.0 kV) and mass spectra acquired over a range of 50 to 2,000 m/z on a G2-Si HDMS mass spectrometer with ion mobility separation (Waters) in enhanced high definition MS (HDMS^E^) or high definition MS (HDMS) mode for non-deuterated and deuterated samples, respectively (*38*). [Glu1]-Fibrinopeptide B standard (Waters) was employed for lock mass correction. During each run, the pepsin column was washed three times with 80 µl of 4% (v/v) acetonitrile and 0.5 M guanidine hydrochloride and blanks were performed between each sample. Three technical replicates (independent H/D exchange reactions) were measured per incubation time.

Peptides were identified with the software ProteinLynx Global SERVER (PLGS, Waters) from the non-deuterated samples acquired with HDMS^E^ by employing low energy, elevated energy and intensity thresholds of 300, 100 and 1,000 counts, respectively. The identified ions were matched to peptides with a database containing the amino acid sequences of the respective proteins, porcine pepsin and their reversed sequences with the following search parameters: peptide tolerance = automatic; fragment tolerance = automatic; min fragment ion matches per peptide = 1; min fragment ion matches per protein = 7; min peptide matches per protein = 3; maximum hits to return = 20; maximum protein mass = 250,000; primary digest reagent = non-specific; missed cleavages = 0; false discovery rate = 100. Deuterium incorporation into peptides was quantified with DynamX 3.0 software (Waters). Only peptides that were identified in all non-deuterated samples and with a minimum intensity of 10,000 counts, a maximum length of 30 amino acids, a minimum number of two products, a maximum mass error of 25 ppm and retention time tolerance of 0.5 min were considered for analysis. All spectra were manually inspected and, if necessary, peptides omitted (e.g., in case of low signal-to-noise ratio or presence of overlapping peptides).

### 4.10 Thermal shift assay

20 µM sVP40_Δ43_ WT/L117A/W95A were mixed with SYPRO Orange (50x final concentration) and VP40 buffer (25 mM Tris, 300 mM NaCl, pH 8) ad 20 µL. A melting curve analysis using a StepOne™ Real-Time PCR system, with a temperature range from 25 °C to 99 °C with a continuous increase of 0.05 °C/sec was performed. Each reaction was measured in triplicates and 20 µL were used per well of a 96 well plate. Negative controls included samples without protein.

## 5 Acknowledgements

The authors would like to thank the BSL4 team Dr. Markus Eickmann, Gotthard Ludwig and Sebastian Schmidt as well as Astrid Herwig, Katharina Kowalski and Nils Krapoth for excellent technical assistance. Also, we would like to thank Dr. Viktoria Reithofer, Dr. Lukas Korf and Dr. Hans-Joachim Emmerich of the Essen lab in Marburg for the kind support regarding crystal data collection at the Swiss Light Source of the Paul-Scherrer Institute in Villigen, Switzerland. Support by the MarXtal facility in Marburg, Germany, for the set-up of crystallization plates, especially Ralf Pöschke, is gratefully acknowledged, as well as support by the Bange lab, University of Marburg, for the kind permission to use their Microfluidizer. The authors would also like to thank Prof. Markus Schnare (University of Marburg) for the support regarding the SEC. This project is funded by the State of Hesse, LOEWE Center “DRUID”, project A1 as well as the “UNISCIENTIA STIFTUNG VADUZ”. We thank the DFG for support through the Marburg core facility for Interaction, Dynamics and Assembly of Biomolecular Structures (Project numbers 260989694 and 324652314).

## 6 Author contributions

Conceptualization: AW, LW, LOE, SB; Data curation: AW, WS, CV, MS; Formal Analysis: AW, WS, CV, MS; Funding acquisition: AW, SB; Investigation: AW, WS, CV, MS; Project administration: AW, SB; Visualization: AW, WS; Writing – original draft: AW, SB; Writing – review & editing: AW, WS, CV, MS, LW, GB, LOE, SB.

## 8 Inclusion and diversity

One or more of the authors of this paper self-identifies as a gender minority in their field of research. We support inclusive, diverse, and equitable conduct of research.

## 9 Supplemental material

### 9.1 Supplemental figures

**Figure S1:**
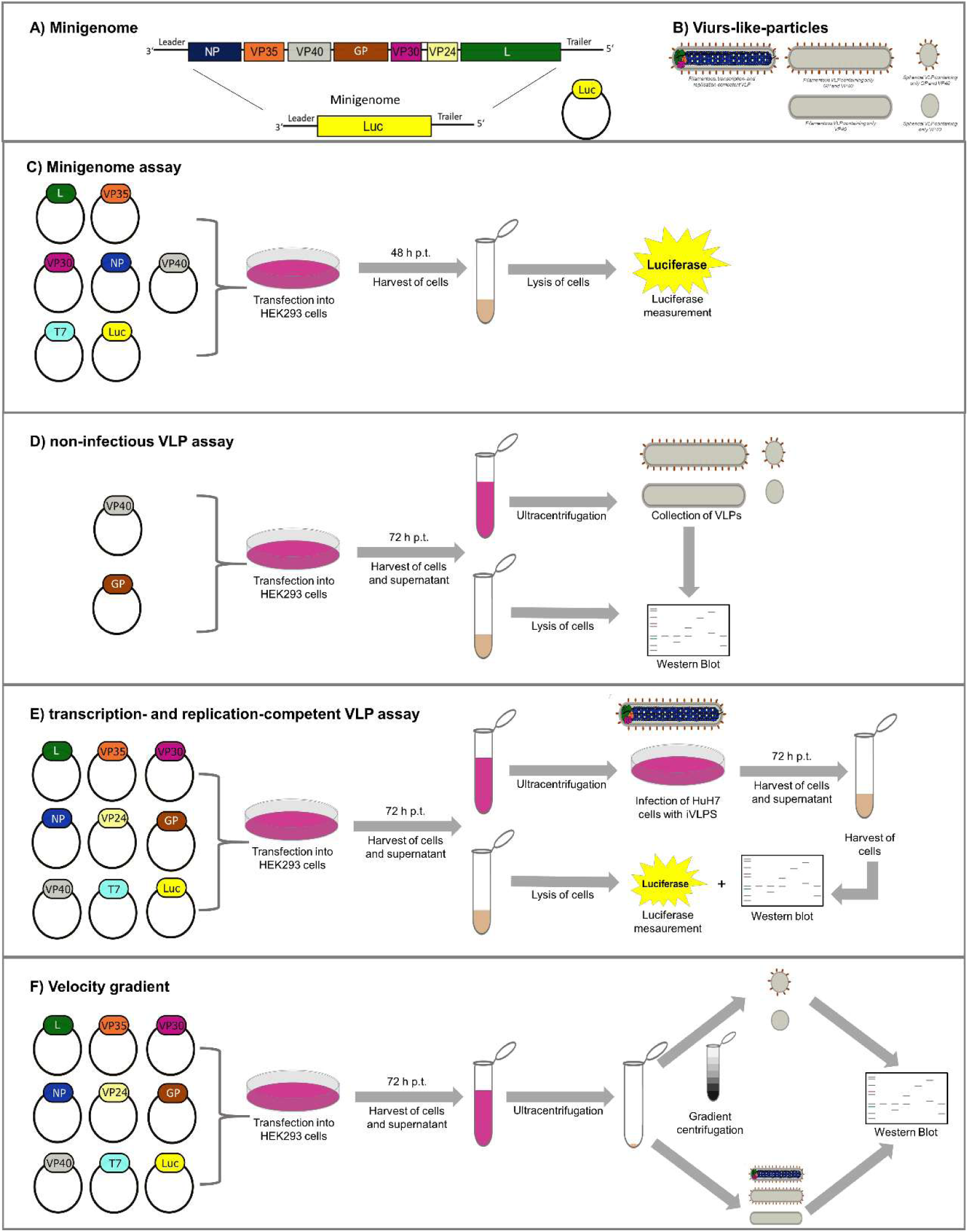
Schematics of minigenome (MG), (transcription- and replication-competent) VLPs (and velocity gradient experiments.

**Figure S2:**
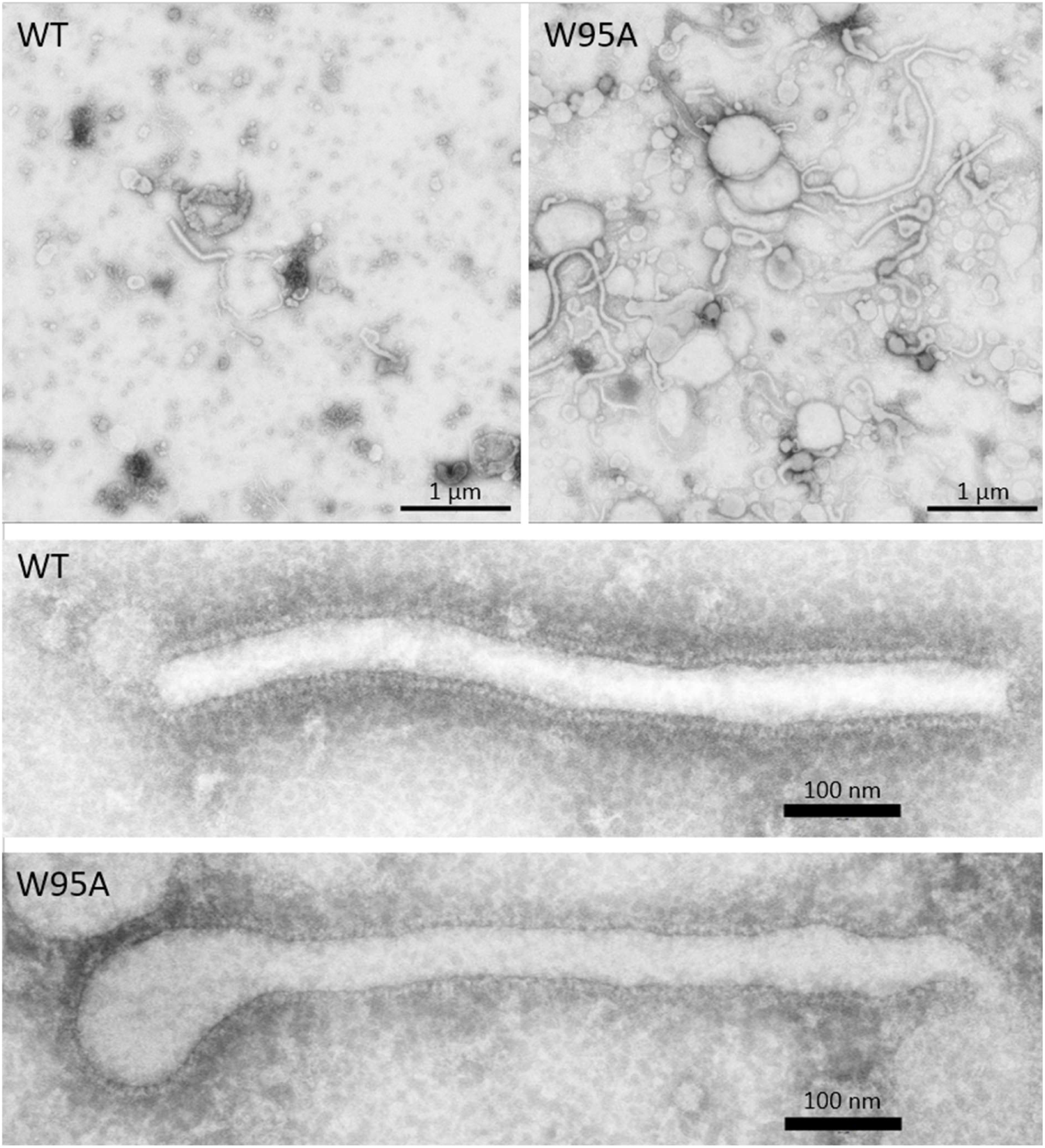
WT and W95A VLP preparations contain morphologically similar filamentous VLPs that are decorated with GP. VLPs were negatively stained with 2% (w/v) phosphotungstic acid and observed in a transmission electron microscope.

**Figure S3:**
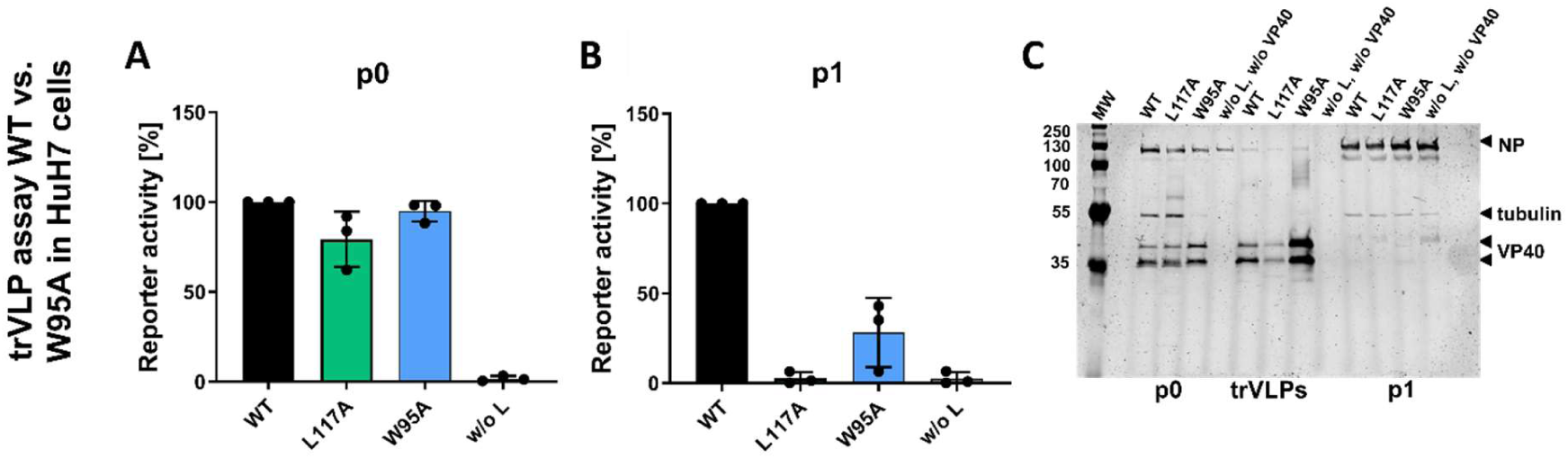
sVP40 W95A results in less trVLPs released from p0 HuH7 cells compared to the WT. P0 producer cells (HuH7) were transfected with pCAGGS NP, VP30, L, VP35, pANDY EBOV minigenome (3E5E), pCAGGS T7, pGL4 firefly luciferase, and either pCAGGS sVP40 WT or W95A. P0 cells were lysed 72 h p.t., and supernatants cleared of cellular debris. trVLPs were collected by ultracentrifugation (2 h at 40,000 rpm) and resuspended in 100 µL PBS_def_ 30 µL of trVLPs were analyzed by WB; 70 µL were used to infect pre-transfected P1 cells. At 72 h post infection (p.i.), p1 cells were lysed and reporter gene activity measured (A for p0 and B for p1 cells). C) WB analyses of p0 and p1 lysates were performed using chicken α-NP, rabbit α-VP40, and mouse α-tubulin as primary antibodies, with donkey α-mouse and goat α-rabbit IRDye® 680 as secondary antibodies. Bars represent mean ± SD of at least three independent experiments.

**Figure S4:**
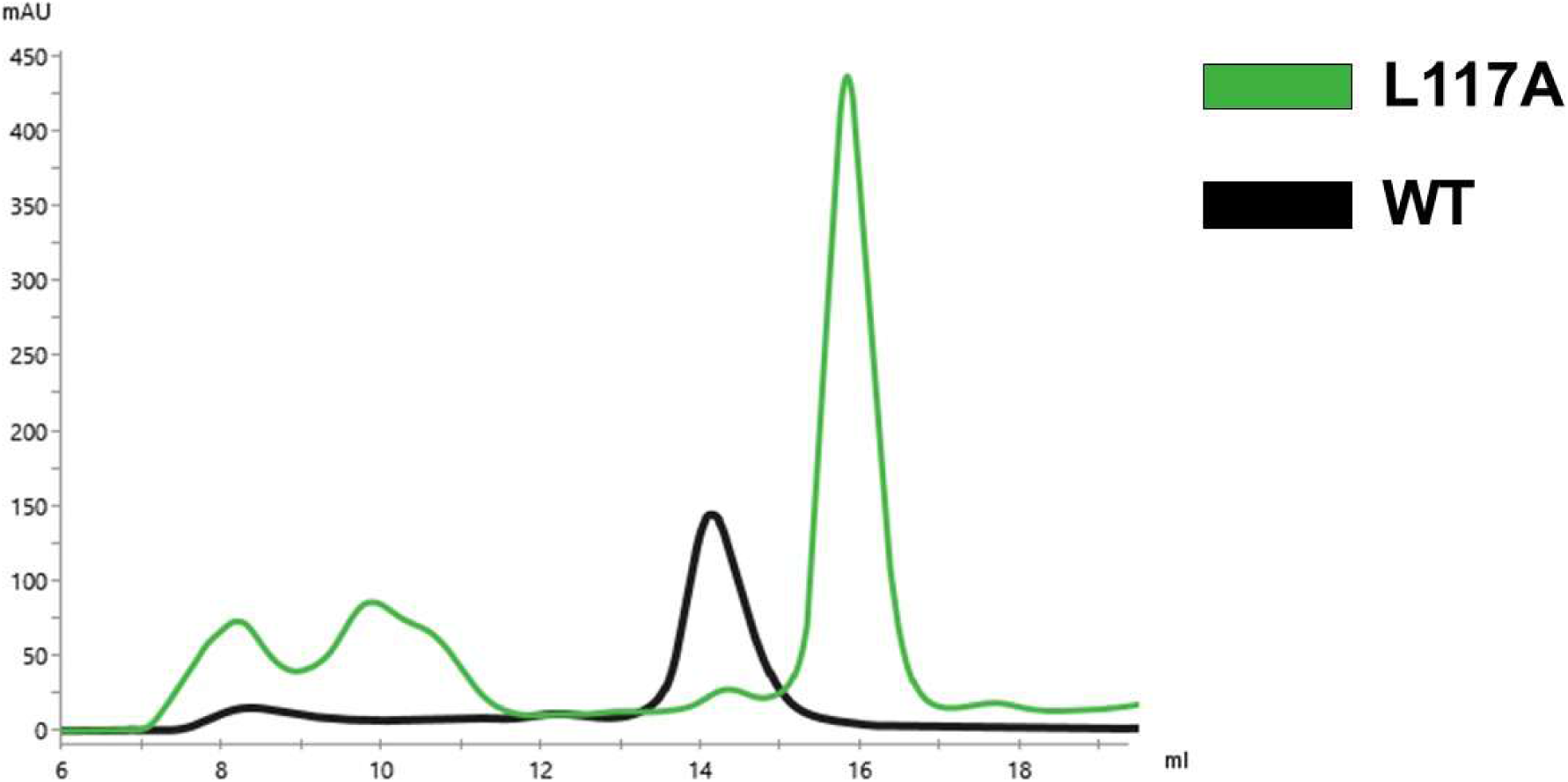
Size-exclusion chromatography profile of sVP40 L117A. A) Size-exclusion chromatography elution profiles of sVP40_Δ43_ WT (black) and sVP40_Δ43_ L117A (green) using a Superdex 200 Increase column (Cytiva).

**Figure S5:**
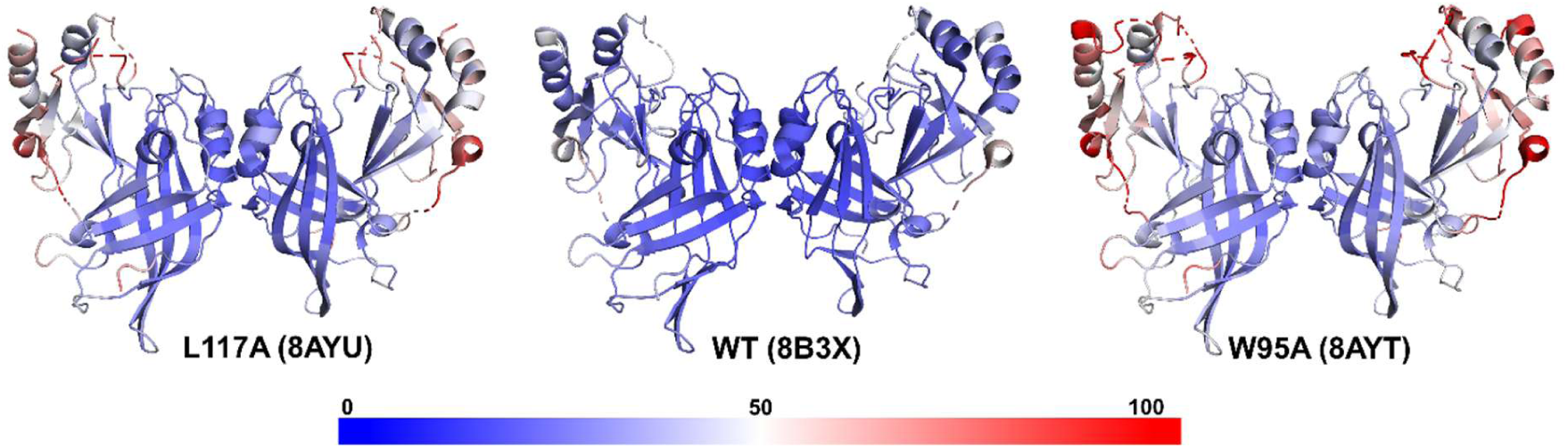
Crystal structures colored according to B-factors with dark blue indicating low levels and dark red indicating high levels.

**Figure S6:**
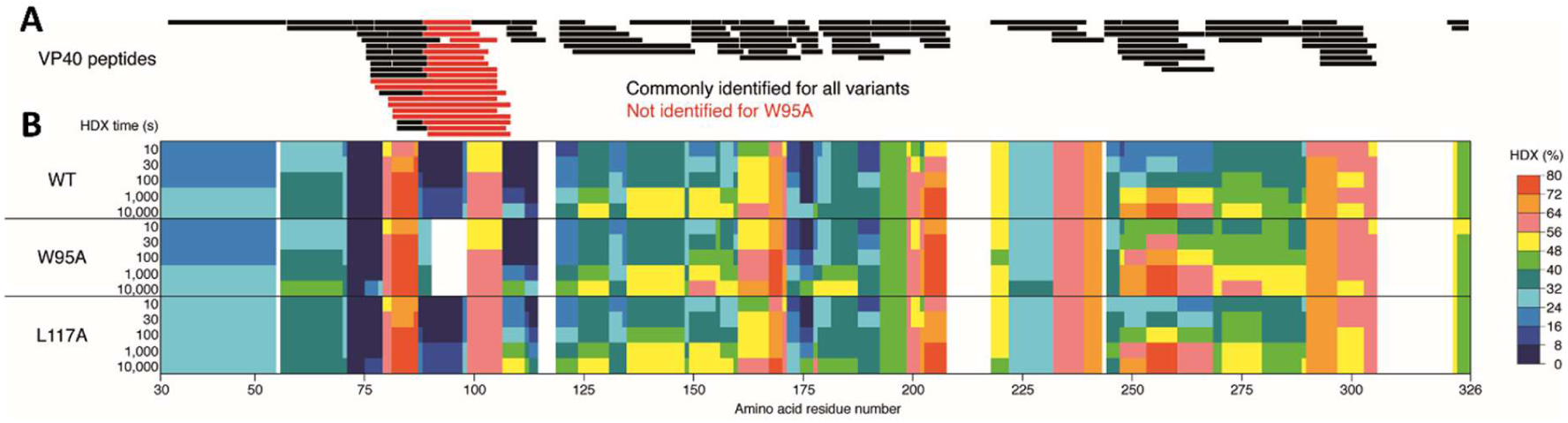
Conformational changes of VP40 variants by HDX-MS. A) VP40 peptides identified in HDX-MS experiments are indicated as black bars and displayed along the amino acid sequence of VP40. The degree of HDX for VP40 WT and its variants is shown in rainbow colors from 0-80% (B).

**Figure S7:**
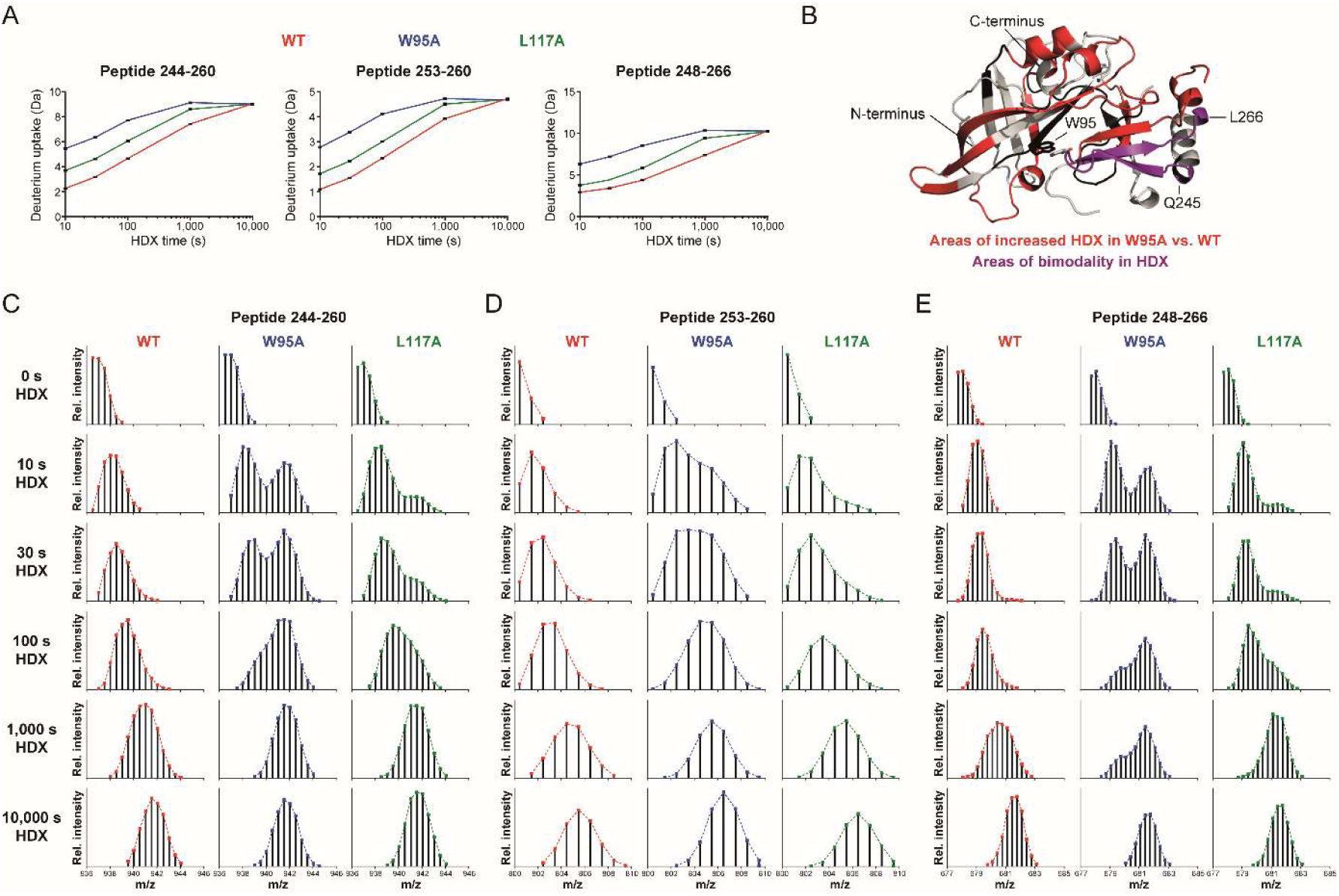
Impact of variation on VP40 conformational flexibility. A-B) Location of selected peptides exhibiting bimodality in HDX displayed on the crystal structure of VP40 (PDB-ID 8B2U (*30*)). C-E) Mass spectra of three representative VP40 peptides exhibiting mixed EX1/EX2 kinetics for hydrogen/deuterium exchange, i.e., C) the peptides containing residues 244-260, D) residues 253-260, and E) residues 248-266. For the L117A and in particular more for the W95A variant, a fast-exchanging population is apparent, which is indicative for at least partial unfolding of secondary structures.

**Figure S8:**
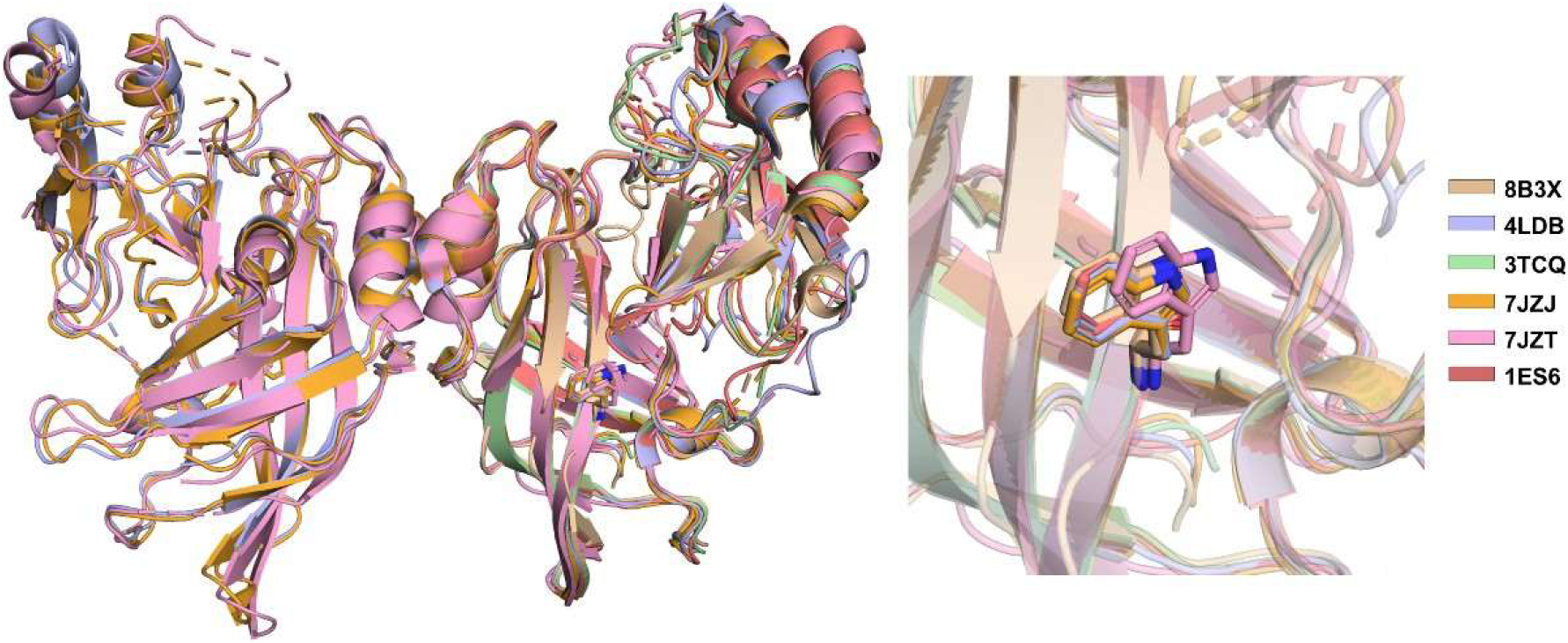
Position and orientation of residue W95 in different VP40 structures. Alignment of dimeric sVP40 (8B3X (*14*) and 3TCQ (*39*); wheat and green) with monomeric (1ES6 (*40*); red) and dimeric eVP40 (4LDB (*3*), 7JZT and 7JZJ (*6*); blue, orange and light pink) with W95 shown as sticks.

**Table S1:** Overview of data obtained by HDX-MS. Each experiment is shown in a separate column.

| Protein | WT | W95A | L117A |
| --- | --- | --- | --- |
| Conditions of HDX | 25 °C in 25 mM Tris-Cl pH 8.0, 300 mM NaCl; final D<sub>2</sub>O = 90 % (v/v) |  |  |
| Time course of HDX | 10/30/100/1,000/10,000 s |  |  |
| Samples | VP40 WT or variants thereof, each at 25 µM stock concentration corresponding to 2.5 µM during HDX |  |  |
| Replicates | 3 technical replicate (separate H/D exchange reactions measured) |  |  |
| Number of Peptides | 115 | 95 | 115 |
| Average peptide length (aa) | 12,6 | 11,37 | 12,6 |
| Sequence coverage (%) | 90,2 | 87,9 | 90,2 |
| Redundancy | 5,41 | 4,14 | 5,41 |
| Back-exchange | No correction for back-exchange based on a fully deuterated sample conducted |  |  |
| Repeatability (average SD) | 0.0179 Da / 0.176% | 0.0213 Da / 0.233% | 0.0229 Da / 0.225% |
| Significance criterium applied | 5% difference in relative HDX |  |  |

### 9.2 Supplemental methods

#### VLPs for negative staining EM

HEK293 cells in a T75 flask (approx. 50% confluency) in 30 ml DMEM with 3% fetal calf serum (FCS) were transfected with 5 µg pCAGGS GP and pCAGGS sVP40 WT/W95A. 48 h p.t. the supernatant was centrifuged for 10 min at 2,500 rpm. After ultracentrifugation 2 h at 4 °C and 40,000 rpm using a 20% sucrose cushion, VLPs were resuspended in 50 µL PBS_def_.

#### Electron microscopy

VLPs were attached to formvar-coated TEM grids and negatively stained with 2% phosphotungstic acid. Electron microscopy was carried out at 120 kV on a JEOL JEM-1400 transmission electron microscope (JEOL, Tokyo, Japan) equipped with a TemCam-F416 camera (TVIPS, Gauting, Germany).

### 9.3 Supplemental dataset

**HDX-MS.** The spreadsheet provides an overview of the conditions of HDX-MS experiments and contains numerical values of HDX observed for VP40 peptides and residues.

